# Small-molecule inhibition of the *Orientia tsutsugamushi* deubiquitylating enzyme OtDUB impairs bacterial reproduction

**DOI:** 10.64898/2026.06.25.734012

**Authors:** Min Jae Lee, Jason R. Hunt, Soyoung Cho, Travis J. Chiarelli, Christofer N. Perry, Jason A. Carlyon, Mark Hochstrasser

## Abstract

Scrub typhus is a potentially fatal infectious disease caused by the obligate intracellular bacterium *Orientia tsutsugamushi*. While antibiotic treatment is generally effective, it requires extended treatment, and drug resistance and treatment failures have emerged. *O. tsutsugamushi* encodes a deubiquitylating enzyme, OtDUB, which interferes with host ubiquitin-dependent pathways. OtDUB cleaves ubiquitin from various substrates, but whether this activity can be selectively targeted by small molecules is unknown. Here we have screened a chemically diverse small-molecule library using a fluorescence-based deubiquitylation assay to identify potential inhibitors of OtDUB. Two compounds, gentisic acid and amiloride hydrochloride, inhibited OtDUB activity at low dosage, with little effect on the related *Wolbachia* CidB or yeast Ulp1 enzymes. Computational docking predicted the compounds engage regions near the OtDUB catalytic pocket, suggesting a competitive mode of inhibition; this was supported by enzyme kinetic analyses. Neither compound caused detectable cytotoxicity in mammalian cells. Amiloride hydrochloride treatment reduced both total cellular deubiquitylating activity and the *O. tsutsugamushi* bacterial load in infected cells. While the identified compounds are not optimized inhibitors, they establish that bacterial pathogen-encoded deubiquitylating enzymes can be targeted by small molecules. Overall, our results provide a framework for using selective inhibitors as tools to study DUB function in genetically intractable intracellular bacteria and as potential treatments for scrub typhus.

## Introduction

Scrub typhus is an acute infectious disease caused by *Orientia tsutsugamushi*, an obligate intracellular bacterium in the order Rickettsiales (Chakraborty & Sarma, 2017). The disease is transmitted to humans through the bites of infected larval *Leptotrombidium* mites (chiggers) and primarily targets endothelial cells, macrophages, and dendritic cells. Scrub typhus has been known since the fourth century to be endemic to the Asia-Pacific region, where it represents one of the most severe rickettsial infections and has an estimated annual incidence exceeding one million cases (Richards & Jiang, 2020; Watt & Parola, 2003; Xu et al., 2017). However, non-travel related cases of scrub typhus, seroprevalence of anti-*Orientia* antibodies, and molecular detection of *Orientia* have also been documented in African, Middle Eastern, and South American countries (Abarca et al., 2020; Bonell et al., 2017; Cosson et al., 2015; Horton et al., 2016; Kocher et al., 2017; Maina et al., 2016; Thiga et al., 2015; Yen et al., 2019). There is potential risk for scrub typhus in North America as *Orientia* spp. DNA was recently detected in trombiculid chiggers in North Carolina, and individuals in this region were determined to be seropositive for antibodies against *Orientia* antigens (Abernathy et al., n.d.; Chen et al., 2023; Richardson et al., 2024). When left untreated, infection can progress to severe complications, including multi-organ dysfunction and death (Vashishtha et al., 2025; Vincent, 2016). The disease is generally responsive to tetracycline-class antibiotics, including doxycycline. Reports of incomplete responses and treatment failures have emerged, however (Srivastava et al., 2024). These observations emphasize the need to explore therapeutic approaches beyond conventional antibiotic treatment.

The ubiquitin–proteasome system (UPS) is a central regulator of protein homeostasis and cellular signaling in eukaryotic cells (Kim et al., 2026; Livneh et al., 2016; Swatek & Komander, 2016). Proteins are targeted for degradation following their modification by ubiquitin, which requires a multistep enzymatic process (Kim et al., 2026). Ubiquitin can be conjugated as a monomer or assembled into polyubiquitin chains linked through distinct lysine residues or its N-terminal methionine, giving rise to functionally diverse signaling outcomes. K48-linked polyubiquitin chains typically target substrates for degradation by the proteasome, whereas other linkages, including K63, play important roles in distinct signaling pathways, particularly those involved in endocytotosis, DNA repair, and immune responses (Finley et al., 1994; Williamson et al., 2013). Ubiquitin can also serve as a covalent signal for autophagy of specific organelles or proteins (Kirkin et al., 2009; Shaid et al., 2013; Yin et al., 2020).

Ubiquitylation is reversible, and deubiquitylating enzymes (DUBs) regulate ubiquitin signaling by processing ubiquitin precursors and removing ubiquitin from modified substrates (Geurink et al., 2019; Hochstrasser, 2025). Eukaryotes encode at least seven different classes of DUBs based on their sequences and mechanisms (Hermanns & Hofmann, 2019). Most are cysteine proteases belonging to the papain superfamily, while a small number are metalloproteases (Clague et al., 2019; Nijman et al., 2005).

Although bacteria lack a canonical UPS, many pathogenic and symbiotic intracellular bacteria encode effector proteins that manipulate host ubiquitin signaling to promote their survival and reproduction. These effectors are delivered into host cells through specialized secretion systems and can function by modifying host ubiquitin pathway components or by acting enzymatically as E3 ligases or DUBs. Most bacterial DUBs identified to date belong to either the OTU (ovarian tumor) family or CE clan of cysteine proteases (Le Negrate et al., 2008; Pruneda et al., 2016). In eukaryotes, CE clan proteases include ubiquitin-like protein-specific proteases (ULPs) that process ubiquitin-like modifiers such as SUMO and NEDD8 (S.-J. Li & Hochstrasser, 1999; Ronau et al., 2016). Bacterial CE clan effectors share the conserved catalytic architecture but have evolved distinct substrate specificities, generally acting preferentially on ubiquitin-linked substrates (Ronau et al., 2016; Pruneda et al., 2016).

A deubiquitylating enzyme from *O. tsutsugamushi*, named OtDUB, was recently identified and structurally characterized (Berk et al., 2020; Lim et al., 2020). The ∼154 kD OtDUB protein belongs to the ULP1-like CE clan proteases; it has a conserved His–Asp–Cys catalytic triad and exhibits DUB activity toward long ubiquitin chains of multiple linkage types, including K48- and K63-linked chains (Berk et al., 2020). Structural analyses showed that OtDUB engages multiple ubiquitin molecules during substrate recognition, including binding through a high-affinity ubiquitin-binding domain (UBD) just downstream of the N-terminal DUB domain (Lim et al., 2020). In human tissue culture cells, OtDUB has been shown to interfere with multiple host membrane-trafficking mechanisms (Berk et al., 2022; Adcox et al., 2022). Homologous proteins with similar deubiquitylating activity have been identified in other pathogenic *Orientia* species, suggesting the conserved role for OtDUB in *Orientia* biology and pathogenesis (Berk et al., 2020; Izzard et al., 2010). These features position OtDUB as a potential target for host-directed therapeutic intervention. Gene mutation or overexpression in *Orientia* bacteria has not yet been achieved, so a chemical-biology approach to assess OtDUB contributions to the *Orientia* life cycle could be a valuable alternative.

Despite the potential importance of OtDUB in *Orientia* infection of human cells (Adcox et al., 2022), no small-molecule inhibitors targeting the enzyme have been reported. In the present study, we sought to identify chemical inhibitors of OtDUB using a high-throughput screening approach. We screened a chemically diverse small-molecule library using a fluorescence-based deubiquitylation assay and identified gentisic acid and amiloride hydrochloride as candidate inhibitors. We further evaluated the specific inhibitory activity of these compounds against purified OtDUB and related ULP1-like proteases, computational docking, kinetic studies, and mammalian cell-based infection assays. Our findings provide the first evidence that OtDUB activity can be chemically inhibited and establish a framework for the development of OtDUB-targeted strategies against scrub typhus.

## Results

### High-throughput screening identifies small-molecule inhibitors of OtDUB

To enable identification of chemical inhibitors of OtDUB, we established a biochemical assay to measure OtDUB deubiquitylating activity *in vitro*. A 259-residue recombinant OtDUB segment including the N-terminal DUB domain and UBD (OtDUB_1–259_) was expressed in *E. coli* and purified (Supplementary Fig. S1A,B). OtDUB_1–259_ was used for the primary screen and follow-up validation experiments. Deubiquitylating activity was assessed using a fluorogenic ubiquitin–AMC substrate with release of AMC from the C-terminus of ubiquitin monitored by fluorescence. Under the assay conditions used, the fluorescence signal increased linearly over the initial 60 minutes of the reaction, indicating that measurements were performed within the linear range of the enzyme reaction (Supplementary Fig. S1C).

Using this assay, we performed a high-throughput screen of the Gen-Plus library (MicroSource), which consists of 960 structurally diverse, biologically active compounds (Figure 1A). Compounds were provided pre-dispensed in black 384-well plates by the Yale Center for Molecular Discovery using automated liquid handling, with 40 nL of each compound per well to achieve a final screening concentration of 20 µM. Fluorescence signals were measured after 60 minutes of reaction and normalized to vehicle-treated controls on each plate to account for plate-to-plate variability. Across screening plates, assay performance remained consistent, allowing comparison of inhibitory effects between compounds. For the high-throughput screen, Z′ factor values were calculated for each plate and ranged from 0.73 to 0.91 (notably, values between 0.5 and 1 were consistently obtained), indicating that assay performance was suitable for screening (Zhang et al., 1999).

**Figure 1.**
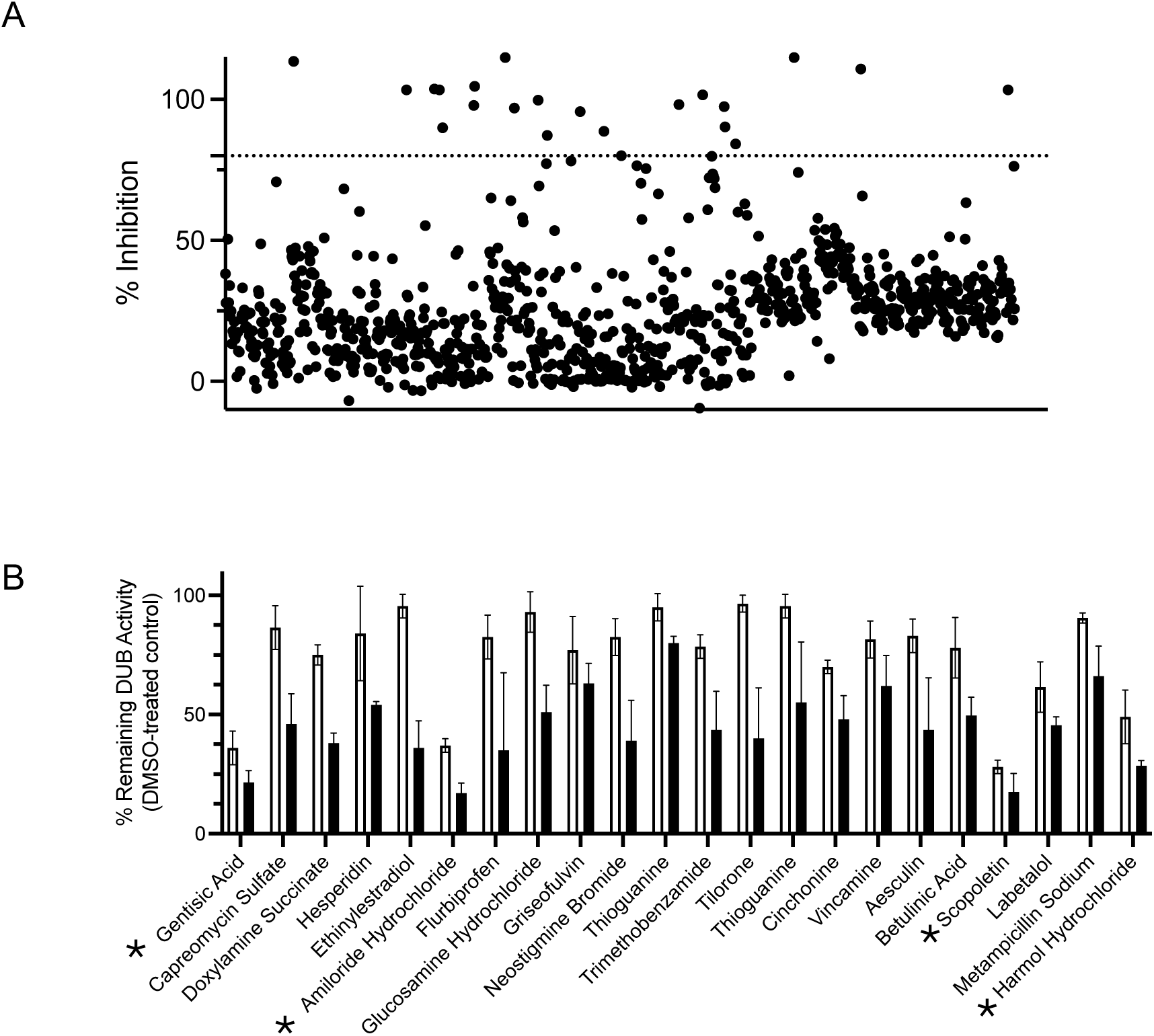
Primary screen for candidate inhibitors of OtDUB activity. (A) A fluorescence-based deubiquitylation assay was used to screen the MicroSource Gen-Plus bioactive compound library for inhibitors of OtDUB activity. Compounds were dispensed at a final concentration of 20 µM and incubated with 125 pM purified OtDUB_1–259_ for 1h at room temperature before initiation of the reaction with the fluorogenic ubiquitin–AMC substrate. The same enzyme concentration and preincubation conditions were used for the secondary validation assays shown in panel B except that compounds were tested at 10 and 20 µM. Deubiquitylating activity was quantified by measuring fluorescence from free AMC after 60 minutes at room temperature. Inhibition was calculated as percent inhibition relative to DMSO-treated controls. Each data point represents an individual compound tested. The dashed line indicates the cutoff used to define preliminary hit compounds (> 80% inhibition), yielding 21 candidate compounds that were selected for secondary validation. In a small number of wells, calculated inhibition values exceeded 100% because fluorescence signals fell below the DMSO-normalized baseline. These values were considered complete inhibition for hit selection and were advanced to secondary validation. (B) Top hits from the primary screen were subjected to secondary validation using a fluorescence-based deubiquitylation assay. The 21 compounds were retested at two concentrations (10 and 20 µM) against purified OtDUB_1–259_. Data shown represent duplicate measurements; error bars indicate the range between duplicate values. Stars indicate the four compounds selected for further characterization.

From the primary screen, compounds that inhibited more than 80% of OtDUB activity relative to vehicle-treated controls were designated as preliminary hits. This cutoff yielded 21 candidate compounds (Supplementary Table 1). Because the primary screen was performed at a single compound concentration, these hits were subjected to secondary validation at both 10 and 20 µM concentrations to assess reproducibility and to exclude false positives (Figure 1B) (Baell & Holloway, 2010; Simeonov et al., 2008; Thorne et al., 2010).

The secondary screen allowed direct comparison of inhibitory activity across several compound concentrations. Compounds were considered reproducible inhibitors if they consistently reduced OtDUB activity by more than 50% relative to vehicle-treated controls across replicate experiments. During secondary validation, the majority of the initial hit compounds failed to reproducibly inhibit OtDUB activity. Many compounds showed minimal inhibition at 10 µM and only modest effects at 20 µM, while others displayed variable responses between the replicates or elevated background signals, consistent with assay interference rather than specific enzymatic inhibition. In contrast, gentisic acid, amiloride hydrochloride, harmol hydrochloride, and scopoletin consistently inhibited OtDUB activity across replicate experiments (Figure 1B). At 10 µM, these compounds reduced OtDUB activity to approximately 26–57% of vehicle-treated controls, indicating substantial inhibition at the lower concentration. Increasing the compound concentration to 20 µM resulted in further suppression of OtDUB activity, with remaining activity reduced to ∼12–30% for these compounds.

Based on their reproducible inhibition at both concentrations, these four compounds were selected for further characterization. Retesting using freshly prepared compound stocks yielded comparable level of inhibition, supporting the robustness of the observed effects and arguing against compound degradation or handling-related artifacts. To confirm that inhibition by these four compounds was not dependent on the specific OtDUB construct used in the primary assays, inhibition experiments were repeated using a shorter OtDUB fragment (OtDUB_1–177_) that lacked the high-affinity UBD (K_d_ ∼5 nM) (Berk et al., 2020). All four compounds inhibited OtDUB_1–177_ to a similar extent as the longer OtDUB_1–259_ fragment, with enzymatic activity reduced by ∼62–84% at 20 uM inhibitor concentrations (Supplementary Fig. S1D). These results indicate that inhibition by the selected compounds is not restricted to a particular fragment design and did not depend on high-affinity ubiquitin binding.

### Selective inhibition of OtDUB relative to other CE-clan enzymes

To determine whether the four prioritized compounds inhibit OtDUB selectively or act more broadly on related cysteine proteases, we compared their activity against OtDUB and several other CE-clan members. For this comparison, we selected the DUB domain of CidB*^w^*^Pip^ (Beckmann et al., 2017; Wang et al., 2022) and the C-terminal catalytic domain of yeast SUMO-specific Ulp1 (Ulp1-C275) (S. J. Li & Hochstrasser, 1999; Mossessova & Lima, 2000), both of which belong to the Ulp1-like CE clan and share a conserved catalytic fold with OtDUB (Berk et al., 2020), while differing in overall sequence and substrate specificity. The two comparator enzymes provide a relevant reference for assessing selectivity within the CE clan (Beckmann et al., 2017; Pruneda et al., 2016, 2018).

Recombinant CidB DUB domain and Ulp1-C275 were purified, and cleavage activities were measured using ubiquitin–AMC and SUMO1-AMC, respectively, under the same reaction conditions employed for OtDUB (Figure 2A, B; Supplementary Fig. S2A). Gentisic acid and amiloride hydrochloride inhibited OtDUB in a concentration-dependent manner (Figure 2C, D). Nonlinear regression analysis of the dose–response curves yielded estimated IC₅₀ values of approximately 12.3 µM for gentisic acid and 8.7 µM for amiloride hydrochloride. By contrast, inhibition of the CidB DUB and Ulp1-C275 required substantially higher compound concentrations, and dose–response curves for these enzymes revealed a shift to higher apparent IC₅₀ values relative to those obtained for OtDUB. At compound concentrations near the OtDUB IC₅₀, only minimal inhibition of CidB and Ulp1-C275 was detected, indicating a window of preferential OtDUB inhibition. Only at higher concentrations did inhibition of the comparator enzymes become apparent. These dose–response relationships further support the conclusion that gentisic acid and amiloride hydrochloride preferentially target OtDUB over other CE-clan proteases. For the remaining two candidate OtDUB inhibitors, harmol hydrochloride and scopoletin, inhibition of OtDUB, CidB, and Ulp1-C275 was similar, consistent with broader activity across CE-clan enzymes (Supplementary Figure 2B). Based on these selectivity profiles, gentisic acid and amiloride hydrochloride were prioritized for subsequent mechanistic and cellular analyses.

**Figure 2.**
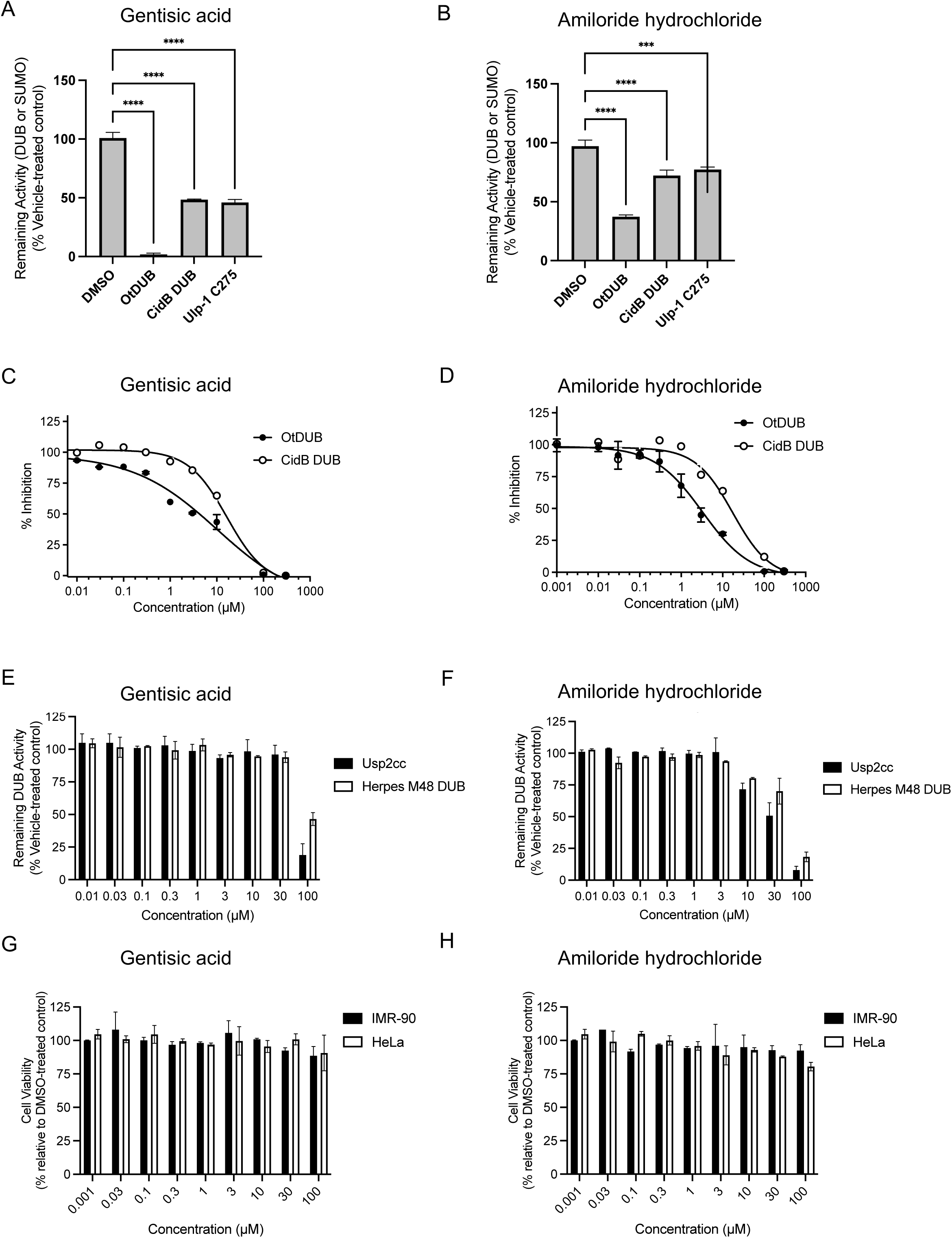
Selectivity profiling of OtDUB inhibitors across diverse USP and CE clan enzymes. (A, B) Protease activity of purified OtDUB_1–259_, the DUB domain of *Wolbachia* CidB*^w^*^Pip^, and the C-terminal catalytic domain of yeast Ulp1 (Ulp1-C275) were measured using fluorogenic AMC-conjugated substrates. OtDUB_1–259_ and CidB^wPip^ DUB were assayed with 200 nM ubiquitin–AMC, whereas Ulp1-C275 was assayed with 1 µM SUMO1-AMC. Reactions contained 125 pM OtDUB_1–259_, 2 nM CidB^wPip^ DUB, or 1 nM Ulp1-C275 and were monitored by fluorescence over time. Data represent mean ± SD from three technical replicates. (C, D) Dose-dependent inhibition of purified OtDUB_1–259_ and *Wolbachia* CidB DUB by selected compounds. Gentisic acid (C) and amiloride hydrochloride (D) were tested over the indicated concentration range by the ubiquitin–AMC fluorescence assay. Enzyme activity was normalized to DMSO-treated controls and plotted as percent inhibition. Data represent mean ± SD from three technical replicates. Dose–response curves were fit by nonlinear regression in GraphPad Prism. (E, F) Dose-dependent effects of gentisic acid and amiloride hydrochloride on structurally distinct DUBs from other families. Gentisic acid (E) and amiloride hydrochloride (F) were tested against the human USP2 catalytic core (USP2cc) and the herpesvirus-encoded M48 DUB over the indicated concentration range using the ubiquitin–AMC cleavage assay. Enzymatic activity was normalized to DMSO-treated controls and plotted as percent inhibition. Data represent mean ± SD from three technical replicates. Dose–response curves were fit by nonlinear regression in GraphPad Prism. (G, H) Assessment of compound cytotoxicity in human cells. IMR-90 fibroblasts and HeLa cells were treated with increasing concentrations of gentisic acid (G) and amiloride hydrochloride (H), and cell viability was measured using a CellTiter-Glo luminescent assay. Data represent mean ± SD from three technical replicates.

Gentisic acid (2,5-dihydroxybenzoic acid) is a small phenolic acid and a known metabolite of salicylates, including aspirin, and has been reported to exhibit diverse biological activities in non-ubiquitin-related contexts (Abedi et al., 2020; Clarke et al., 1953; Hosseinzadeh et al., 2025). Amiloride hydrochloride is a clinically valuable compound originally developed as a potassium-sparing diuretic and is best characterized as an inhibitor of epithelial sodium channels, with additional reported activities at higher concentrations (Bull & Laragh, 1968; Sun & Sever, 2020; Vidt, 1981). Both compounds are chemically simple, commercially available, and have been characterized in other biological systems (Abedi et al., 2020; Sun & Sever, 2020; Vidt, 1981). These prior characterizations reduce concerns related to compound stability and nonspecific assay interference.

### Evaluation of compound specificity against other DUBs and cytotoxicity in human cells

To examine whether gentisic acid and amiloride hydrochloride inhibit DUB enzymes beyond the CE clan, we tested their activity against structurally distinct DUBs from other families. As representative examples, we selected the human USP2 catalytic core (USP2cc), a representative USP-family DUB and the herpesvirus-encoded M48 DUB, a viral tegument protein–associated DUB. Both enzymes are unrelated to Ulp1-like CE-clan enzymes except that they share a papain-like cysteine protease fold (Renatus et al., 2006; Schlieker et al., 2007). Purified recombinant USP2cc and M48 were assayed using established assay conditions across a concentration range extending up to 100 µM (Figure 2E and 2F). Under these conditions, gentisic acid and amiloride hydrochloride caused little if any inhibition of either USP2cc or M48 except at the highest concentrations of inhibitor that are well above the range required to inhibit OtDUB. These observations indicate that these compounds do not act as broad-spectrum DUB inhibitors and instead show a bias toward inhibition of OtDUB and closely related Ulp1-like enzymes.

We next evaluated the cytotoxic effects of gentisic acid and amiloride hydrochloride in human cells. Normal human IMR-90 fibroblasts and HeLa cells were treated with increasing concentrations of each compound, and cell viability was assessed using a luminescent CellTiter-Glo assay. Neither compound induced detectable cytotoxicity at concentrations up to 100 µM in either cell line (Figure 2G and 2H). These results indicate that substantial chemical inhibition of OtDUB can be achieved without causing detectable toxicity to host human cells under the conditions tested.

### Inhibition of OtDUB activity in human cells overexpressing OtDUB

Having established that gentisic acid and amiloride hydrochloride do not broadly inhibit members of other DUB families and are well tolerated by human cells, we next examined whether these compounds inhibit OtDUB activity in a cellular context. To examine this, HeLa cells were transiently transfected with a plasmid encoding full-length OtDUB fused to an N-terminal 3xFLAG-6His tag (Lim et al., 2020). Twenty-four hours after transfection, cells were treated with gentisic acid or amiloride hydrochloride for an additional 24 hours prior to harvest. Immunoblot analysis confirmed expression of the ∼155 kD FLAG-tagged OtDUB in transfected cells compared to mock-transfected controls (Figure 3A). Deubiquitylating activity was measured in cell lysates using the ubiquitin-AMC substrate; OtDUB overexpression resulted in a ∼80% increase in total DUB activity relative to mock-transfected cells (Figure 3B).

**Figure 3.**
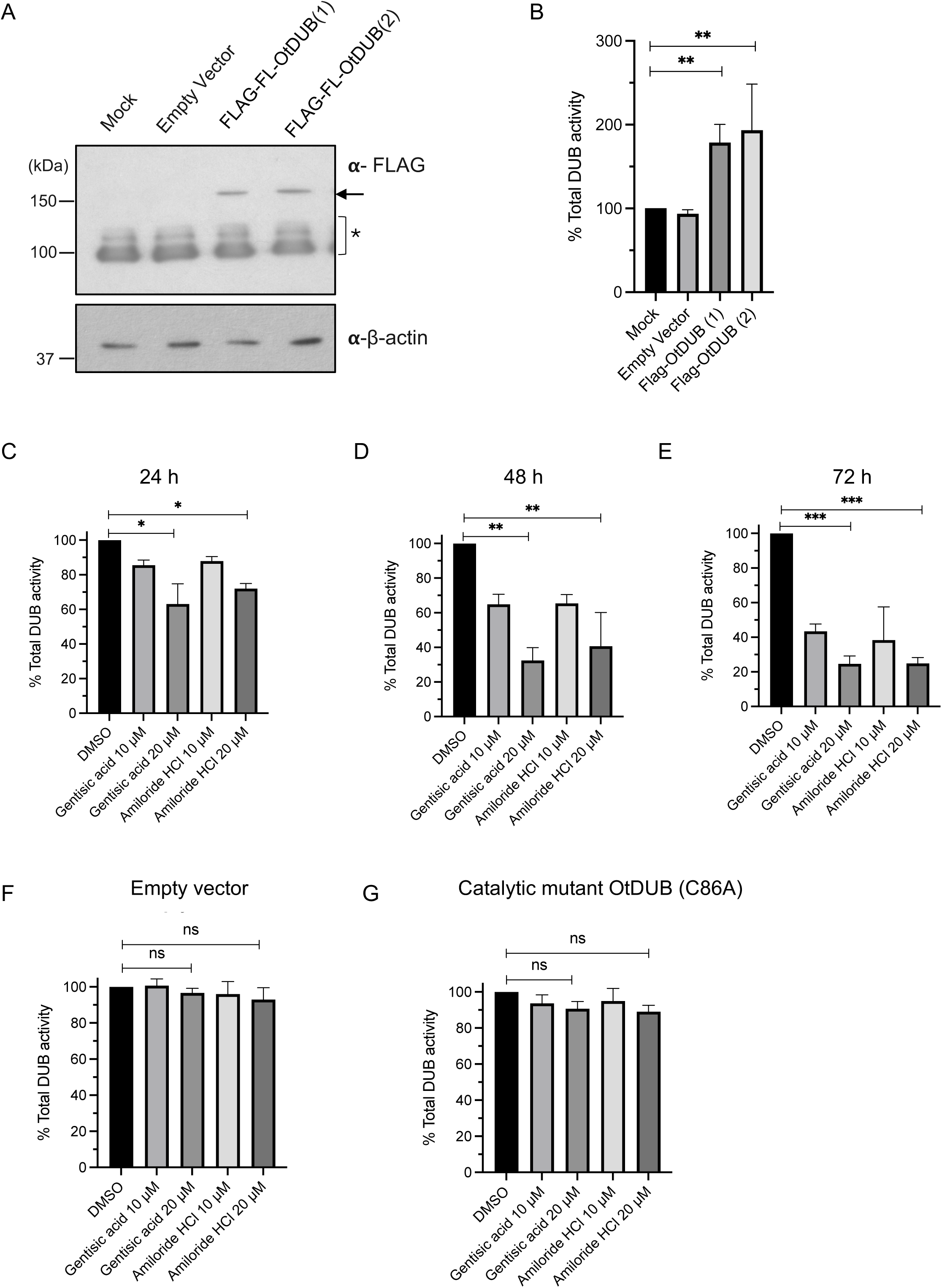
Inhibition of OtDUB activity in human cells overexpressing OtDUB. (A) HeLa cells were transiently transfected with a plasmid encoding full-length FLAG-tagged OtDUB or with empty vector as a control. Cells were harvested 24 h after transfection, and lysates were analyzed by immunoblotting with an anti-FLAG antibody. Full-length FLAG-tagged OtDUB (∼155 kDa) was detected in two independent biological replicates (lanes 3, 4). The arrow indicates the OtDUB band, and the bracketed lower bands marked with an asterisk indicate nonspecific anti-FLAG-reactive bands. β-actin was used as a loading control. (B) Deubiquitylating activity was measured in the same cell lysates from panel A using a fluorogenic ubiquitin–AMC substrate. OtDUB-expressing lysates showed increased total DUB activity compared with mock- or empty vector-transfected controls. DUB activity was normalized to mock-transfected cells. Data are presented as mean ± SEM from three biological replicates. Statistical significance was determined by one-way ANOVA followed by Tukey’s multiple-comparisons test. **P < 0.01. (C-E) Time-dependent effects of gentisic acid and amiloride hydrochloride on DUB activity in OtDUB-overexpressing HeLa cells. HeLa cells transiently expressing full-length FLAG-tagged OtDUB were treated with gentisic acid or amiloride hydrochloride at the indicated concentrations for 24 h (C), 48 h (D), or 72 h (E). Total DUB activity was measured in cell lysates using ubiquitin–AMC fluorescence assays. DUB activity was normalized to DMSO-treated controls at each time point. Data are presented as mean ± SEM from three independent biological replicates. Statistical significance was determined by one-way ANOVA followed by Tukey’s multiple-comparisons test. *P < 0.05, **P < 0.01, ***P < 0.001. (F) Effect of gentisic acid and amiloride hydrochloride on HeLa cells transfected with empty vector after 72 h compound treatment. Data are presented as mean ± SEM from three independent biological replicates. No statistically significant differences were observed relative to DMSO-treated controls as determined by one-way ANOVA followed by Tukey’s multiple-comparisons test. (G) Effect of gentisic acid and amiloride hydrochloride on cells expressing a catalytically inactive OtDUB mutant (C86A). Cells were treated with the indicated concentrations of each compound for 72 h, and DUB activity was measured in cell lysates. Data are presented as mean ± SEM from three independent biological replicates. No statistically significant differences were observed relative to DMSO-treated controls, as determined by one-way ANOVA followed by Tukey’s multiple-comparisons test.

In this expression system, OtDUB contributes substantially to the measured cellular DUB activity, such that specific inhibition of OtDUB should be reflected in a reduction in total DUB activity. Indeed, treatment with gentisic acid or amiloride hydrochloride led to significant reductions in cellular DUB activity in OtDUB-overexpressing cells; the impact was enhanced as treatment time was extended or inhibitor dosage was increased (Figure 3C-E). By contrast, no significant change in total activity was observed in mock-transfected cells under the same treatment conditions for 72 h (Figure 3F).

To determine whether inhibition of relative DUB activity required catalytically active OtDUB, we performed parallel experiments using a catalytically inactive OtDUB mutant (C86A). In cells expressing the mutant protein, treatment with gentisic acid or amiloride hydrochloride failed to reduce total DUB activity relative to the vehicle-treated control (Figure 3G). Immunoblot analysis showed that gentisic acid and amiloride hydrochloride did not reduce the expression level of FLAG-tagged OtDUB over the course of treatment, indicating that the observed decrease in DUB activity was not due to loss of OtDUB protein (Supplementary Figure S3).These results indicate that the DUB inhibition observed in OtDUB-overexpressing cells depends on a catalytically active OtDUB enzyme and is not attributable to nonspecific suppression of cellular DUB activity.

### Docking analysis supports binding near the OtDUB active site

To gain insight into how interaction between OtDUB and the identified compounds leads to enzyme inhibition, *in silico* docking analyses were performed using AutoDock Vina (Eberhardt et al., 2021; Trott & Olson, 2010). Docking simulations were carried out using the available OtDUB structure (Lim et al., 2020), and apparent interactions with the lowest predicted free energy were identified. For gentisic acid, docking yielded a predicted binding mode in which the compound was positioned near the catalytic pocket of OtDUB (Figure 4A). The model placed gentisic acid near Asp96 and Gln129, residues located in the catalytic region. Amiloride hydrochloride was also predicted to dock near the OtDUB catalytic pocket (Figure 4B). These docking models suggest that both compounds can engage regions in or very close to the OtDUB catalytic site.

**Figure 4.**
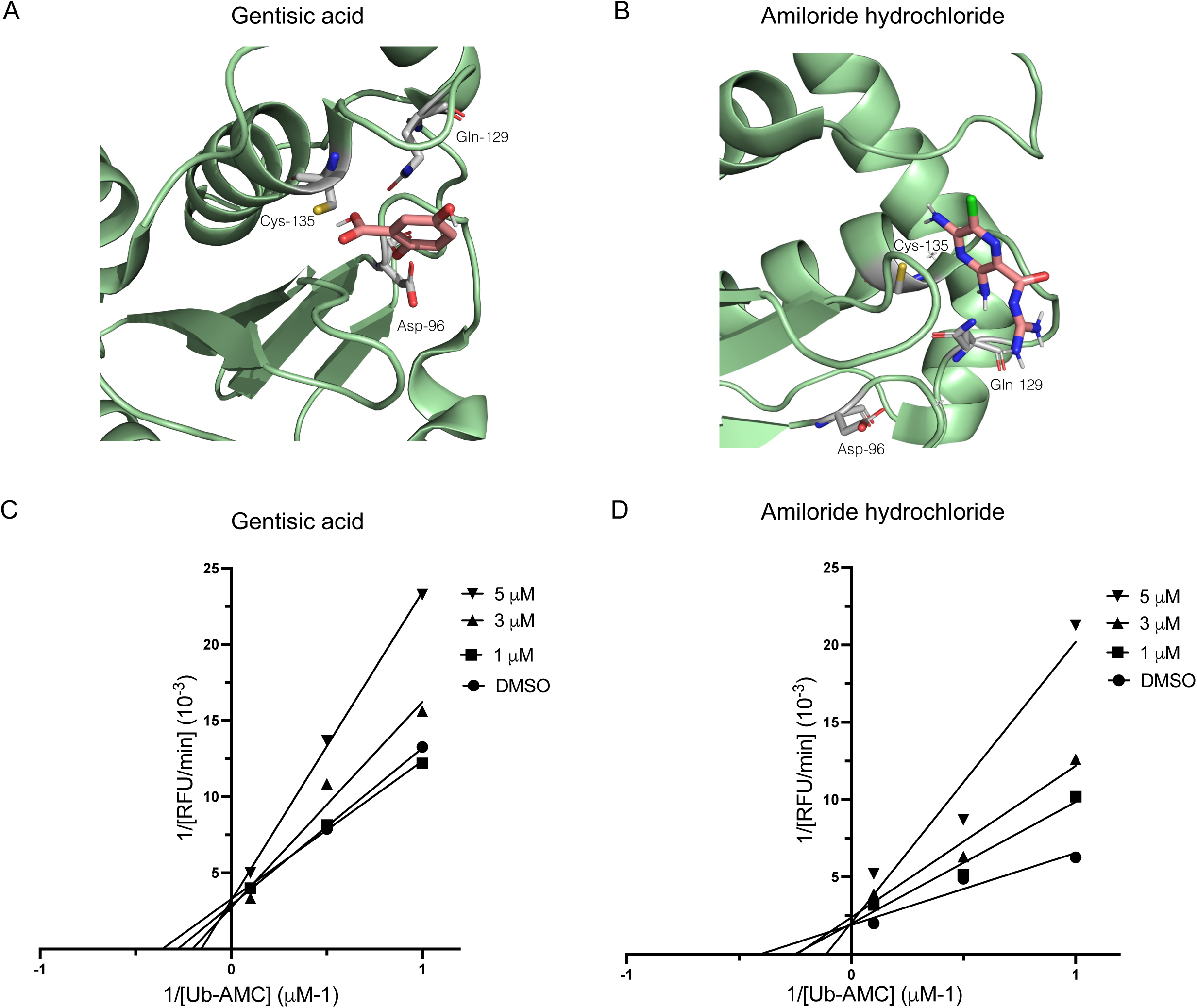
Docking and kinetic analyses suggest competitive inhibition of OtDUB by gentisic acid and amiloride hydrochloride. (A) Predicted binding mode of gentisic acid docked onto the OtDUB deubiquitylase domain (PDB 6UPS). Docking was performed using AutoDock Vina. Gentisic acid is positioned in the catalytic region, with predicted interactions including the highlighted residues. (B) Predicted binding mode of amiloride hydrochloride docked onto OtDUB. The compound is positioned in the catalytic site, with predicted interactions including catalytic triad residues Asp96 and His76. (C) Enzyme kinetics of OtDUB in the presence of gentisic acid. Initial reaction velocities were measured at increasing concentrations of ubiquitin–AMC substrate in the absence or presence of the indicated concentrations of gentisic acid, and data are shown as Lineweaver–Burk plots. (D) Enzyme kinetic analysis of OtDUB activity in the presence of amiloride hydrochloride, analyzed as in (C). For both compounds, the kinetic profiles show changes consistent with competitive inhibition, characterized by shifts in apparent K_m_ with minimal effects on V_max_.

Control docking analyses were performed to compare the docking behavior of gentisic acid and amiloride hydrochloride with unrelated compounds and alternative cysteine proteases. In docking against OtDUB, unrelated compounds including dexamethasone acetate, peruvoside, minocycline hydrochloride, and pomiferin showed variable predicted docking energies and did not reproduce the docking profiles observed for gentisic acid and amiloride hydrochloride (Supplementary Fig. S4A). Gentisic acid and amiloride hydrochloride were also docked against alternative deubiquitylating enzymes, including M48, USP2, SENP1, USP7, USP14, and RickCE. Predicted docking energies varied across these enzymes, including some that were close to OtDUB docking energy, but did not reproduce the sites of interaction observed with OtDUB and amiloride hydrochloride had the most favorable binding energy with OtDUB (Supplementary Fig. S4B).

To examine whether gentisic acid and amiloride hydrochloride inhibit OtDUB through a substrate-competitive mechanism, as predicted from the docking studies, enzyme kinetic analyses were performed using increasing concentrations of ubiquitin–AMC substrate in the presence of different concentrations of each inhibitor (Figure 4C, 4D). For both compounds, increasing inhibitor concentration resulted in an increase in the apparent K_m_ while the apparent V_max_ (y-intercept) remained largely unchanged. These kinetic trends are consistent with a competitive mode of inhibition and suggest that gentisic acid and amiloride hydrochloride interfere with substrate binding to OtDUB.

### Amiloride hydrochloride reduces *O. tsutsugamushi* bacterial burden in host-cell pretreatment assays

To examine whether the compounds identified in the OtDUB screen inhibited deubiquitylating activity in *O. tsutsugamushi*-infected cells and affected bacterial load, we performed cell-based infection assays using HeLa cells, an established model for studying *Orientia*-host cell interactions (Adcox et al., 2024; Allen et al., 2025; Berk et al., 2022; Siff et al., 2026). HeLa cells were treated for 48 h with amiloride hydrochloride or gentisic acid at 25–100 µM or DMSO vehicle control followed by incubation with *O. tsutsugamushi.* As determined by qPCR, amiloride hydrochloride but not gentisic acid significantly reduced the bacterial load and was most effective at a concentration of 100 µM (Supplementary Fig. S5A). Next, we assessed the efficacy of treating host cells for a shorter duration and also distinguished effects on the bacterium from those mediated through the host cell environment. Three treatment regimes were compared: *O. tsutsugamushi* bacteria treated with 100 µM gentisic acid or amiloride hydrochloride for 1 h prior to incubation with untreated HeLa cells; untreated *Orientia* added to HeLa cells that had been treated with either compound for 2 h; or treated bacteria added to treated HeLa cells. Pretreatment of HeLa cells with amiloride hydrochloride but not gentisic acid significantly reduced the *O. tsutsugamushi* load whereas pretreatment of the bacteria had no effect (Supplementary Fig. S5B). In parallel, amiloride hydrocholoride pretreatment of HeLa cells but not *O. tsutsugamushi* measurably reduced total deubiquitylating activity in cell extracts at 24 hpi (Figure 5A). Amiloride hydrochloride treatment of host cells consistently yielded significant inhibition of infection by 48 h post-infection (hpi), inconsistently did so at 24 hpi, and exerted no effect at 2 hpi (Supplementary Fig. S5 and Figure 5B). To determine whether the reduction in bacterial burden reflected impaired bacterial entry, we employed an immunofluorescence microscopy-based assay that differentially immunolabeled extracellular surface-bound versus internalized *Orientia*. Neither gentisic acid nor amiloride hydrochloride detectably inhibited *O. tsutsugamushi* entry (Figure 5C).

**Figure 5.**
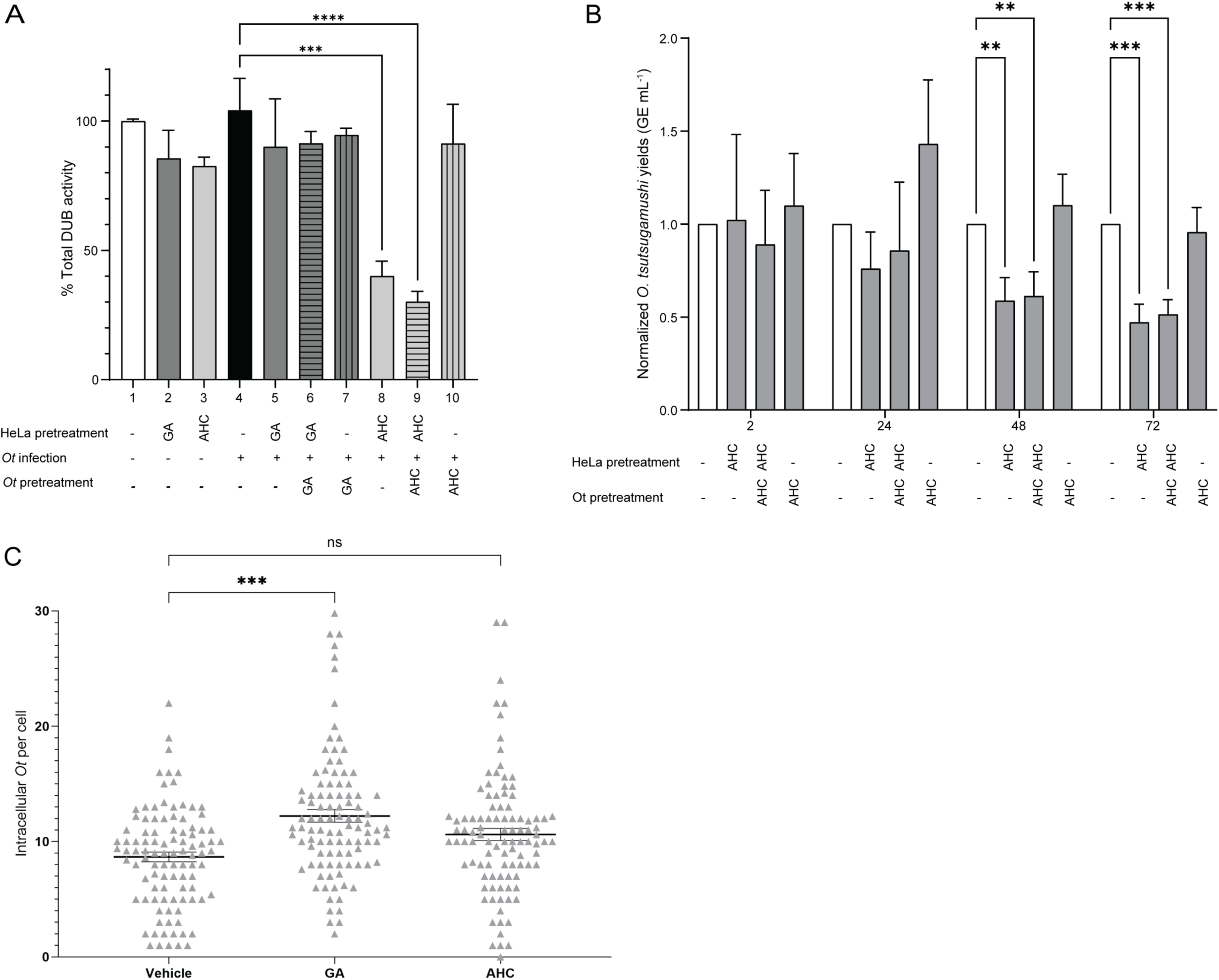
Chemical inhibition of OtDUB is associated with reduced *Orientia* infection in human cells. (A) Total cellular deubiquitylating activity in HeLa cell lysates measured using ubiquitin-AMC. HeLa cells were pretreated with gentisic acid (GA) or amiloride hydrochloride (AHC) prior to infection with *O. tsutsugamushi* as indicated. In selected conditions, host cell-free *O. tsutsugamushi* was pretreated with compound before infection. Total DUB activity was measured in cell lysates and is presented as relative total DUB activity normalized to the uninfected DMSO-treated control (lane 1). Data represent mean ± SD from three biological replicates, each measured in triplicate. Statistical comparisons were performed by one-way ANOVA followed by Dunnett’s multiple-comparisons test using infected DMSO-treated cells (lane 4) as the reference control. ***P < 0.001; ****P < 0.0001. (B) Effect of AHC pretreatment of *O. tsutsugamushi* on infection. Bacteria were pretreated with AHC at 100 µM for 1 h, washed extensively with PBS, and used to infect HeLa cells for 2 h. Alternatively, HeLa cells were treated with AHC at 100 µM for 2 h followed by incubation with treated or untreated *O. tsutsugamushi* bacteria. Following infection, extracellular bacteria were removed by washing and the cells were maintained in compound-free medium. At the indicated post-infection time points, the *O. tsutsugamushi* load measured as GE ml^-1^ was quantified by qPCR. Data represent mean ± SD from three biological replicates, each measured in triplicate. Statistical comparisons were performed by two-way ANOVA followed by Dunnett’s multiple-comparisons test using infected vehicle-treated cells as the reference control. **, *P* < 0.01; ***, *P* < 0.008. (C) AHC and GA fail to inhibit *O. tsutsugamushi* internalization. HeLa cells grown on coverslips and/or *O. tsutsugamushi* bacteria were treated with compounds as in (B), incubated together, and fixed at 2 h. Extracellular bacteria were labeled with membrane-impermeable TSA56 antibody followed by Alexa 488-conjugated secondary antibody. The cells were then permeabilized, and all bacteria (intracellular and extracellular) were immunolabeled with TSA56 antibody followed by Alexa Fluor 594-conjugated secondary antibody. Bacteria that were only Alexa Fluor 594-positive were regarded as internalized, while bacteria stained that were both Alexa Fluor 488- and Alexa Fluor 594-positive were considered extracellular surface-bound bacteria. Data are a composite of five separate experiments. Statistical comparisons were performed by two-way ANOVA followed by Dunnett’s multiple-comparisons test using infected vehicle-treated cells as the reference control. ***P < 0.0001.

We next examined if the reduction in *Orientia* load achieved with amiloride hydrochloride was associated with altered ubiquitylation of the bacterium. *O. tsutsugamushi* organisms were incubated with pretreated HeLa cells and treatment was maintained throughout infection. The cells were fixed, immunolabeled with a pan-ubiquitin antibody, FK2, and antibody against the *O. tsutsugamuhsi* immunodominant TSA56 (56-kDa type-specific antigen) outer membrane protein (Beyer et al., 2017), and co-stained with the DNA dye DAPI. Colocalization of FK2 and TSA56 immunosignals on cytosolic bacteria was scored as *Orientia* being ubiquitylated. As a positive control for FK2 immunolabeling, in parallel we screened cells infected with *Anaplasma phagocytophilum*, a vacuole-adapted rickettsial pathogen that lives in a multivesicular body-like organelle that is ubiquitylated (Huang et al., 2012; Read et al., 2022). In control cells, FK2 immunolabeled 100% of the *A. phagocytophilum* vacuoles but only 6% of cytosolic *O. tsutsugamushi* at 30 min and ≤ 1% at subsequent time points (Fig. 6A, B). While slightly more ubiquitylated *O. tsutsugamushi* bacteria were observed in amiloride hydrochloride treated cells versus controls at all time points, the difference was statistically insignificant. Thus, *O. tsutsugamushi* avoids ubiquitylation and this phenomenon cannot be reversed by amiloride hydrochloride. Overall, these infection assays suggest that the amiloride-associated reduction in *Orientia* burden is most consistent with a host cell-associated effect of OtDUB rather than with direct antibacterial activity or significant impairment of bacterial ubiquitylation.

**Figure 6.**
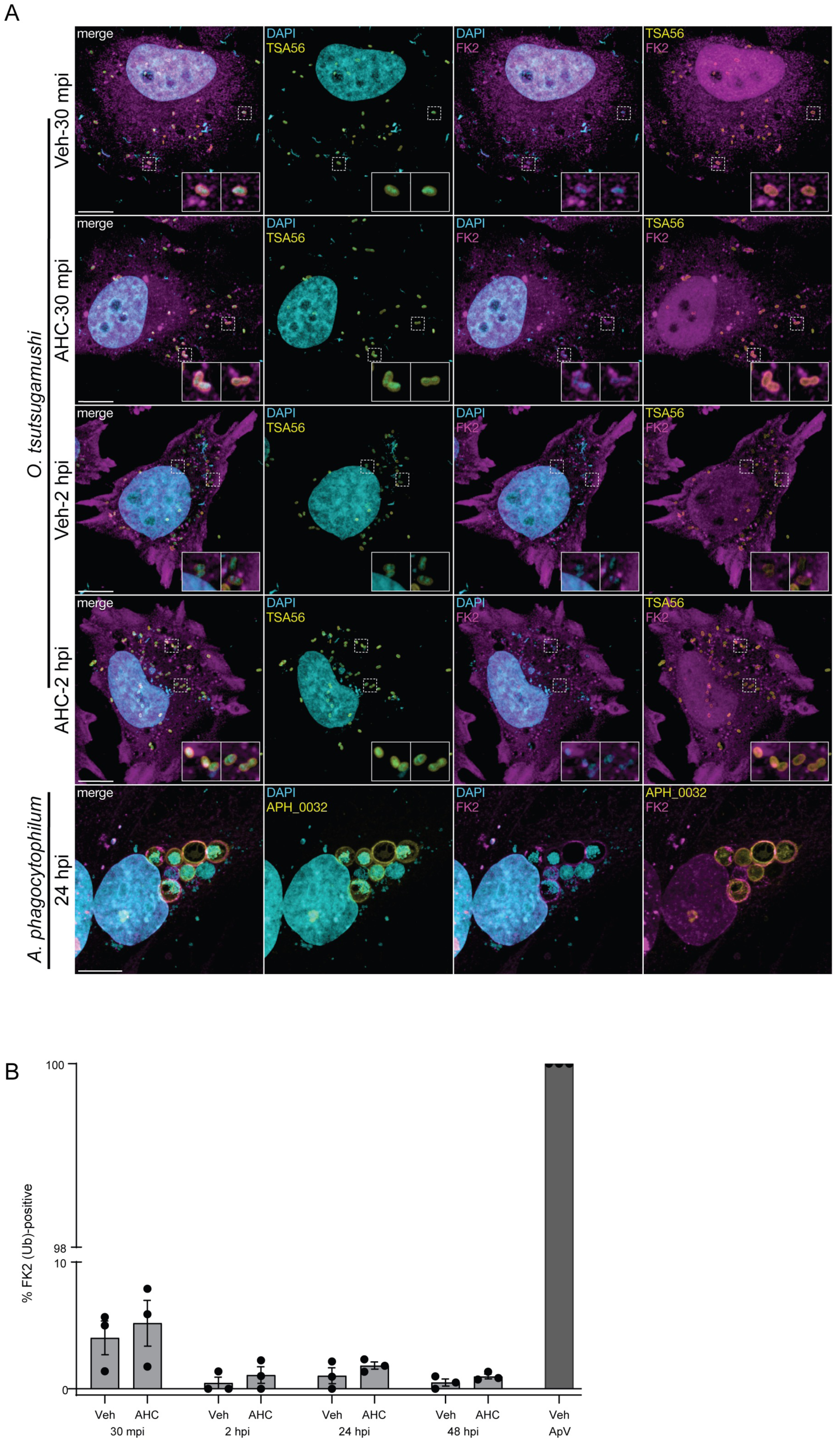
Amiloride hydrochloride does not increase *O. tsutsugamushi* susceptibility to ubiquitylation. (A and B) HeLa cells were treated with amiloride hydrochloride (AHC) or vehicle (Veh) for 2 h followed by incubation with *O. tsutsugamushi*. At 30 min post-infection (30 mpi) or 2 h post-infection (hpi), the cells were fixed, immunolabeled with anti-ubiquitin antibody FK2 and anti-*O. tsutsugamushi* TSA56 (to demarcate individual bacteria), and stained with DAPI (to label host cell nuclei and bacterial nucleoids). As a positive control for FK2 immunolabeling, *A. phagocytophilum* infected RF/6A cells were fixed at 24 hpi, immunolabeled with FK2 and antibody against the *A. phagocytophilum* vacuolar membrane protein APH_0032, and stained with DAPI. (A) Pseudocolored representative maximum intensity Z-projection confocal micrographs of *O. tsutsugamushi*- and *A. phagocytophilum-*infected cells. Regions denoted by hatched lined boxes are magnified (3x) in insets demarcated by solid lined boxes. DAPI channel is log transformed to aid in visualization of bacterial nucleoids. *O. tsutsugamushi* FK2 channel is log transformed. Scale bars, 10 µm. (B) Percentage of FK2-immunolabeled *O. tsutsugamushi* bacteria or *A. phagocytophilum* vacuoles. Fifty to 150 *O. tsutsugamushi* bacteria or *A. phagocytophilum* vacuoles were countered per time point. The percentage of TSA56-positive *O. tsutsugamushi* bacteria or APH_0032-positive *A. phagocytophilum* vacuoles that were also detected by FK2 was determined. Data represent mean ± SD from three biological replicates, each measured in triplicate.

## Discussion

This study addresses whether the *Orientia tsutsugamushi* pathogen-encoded deubiquitylating enzyme OtDUB can be inhibited by small molecules and whether use of such compounds can help define the contribution of OtDUB deubiquitylating activity to pathogen replication. OtDUB engages host ubiquitin signaling through an extended surface that interacts with multiple ubiquitin molecules (Berk et al., 2020; Lim et al., 2020). Although CE-clan deubiquitylating enzymes have been well characterized, it remained unclear whether the CE-clan OtDUB enzyme could be chemically modulated (Luce-Fedrow et al., 2018). Using an unbiased high-throughput screening approach, we identified gentisic acid and amiloride hydrochloride as compounds that reproducibly inhibit OtDUB activity in biochemical assays. Both compounds are well-characterized bioactive molecules, but their ability to inhibit a specific bacterial DUB was not anticipated. Gentisic acid and amiloride hydrochloride are stable in aqueous and cellular environments, reducing concerns related to compound degradation (Mashhadi et al., 2020; Yellepeddi et al., 2020). Prior use of these compounds in unrelated biological contexts also showed that the observed inhibition did not come from assay interference, which is common for poorly characterized screening hits (Baell & Holloway, 2010). Docking analysis and enzyme kinetic measurements provided insight into how these compounds inhibit OtDUB. Computational docking predicted that both compounds bind in or near the catalytic site of OtDUB (Lim et al., 2020). In line with this, kinetic analyses showed a competitive inhibition profile. This mode of inhibition is consistent with direct engagement of the catalytic region and does not require disruption of the broader ubiquitin-binding interface.

In cellular infection assays, amiloride hydrochloride reduced *O. tsutsugamushi* bacterial burden when HeLa cells were pretreated before infection, whereas direct pretreatment of bacteria did not produce a comparable effect. Gentisic acid did not measurably reduce bacterial burden under any tested condition, possibly because of *Orientia*-induced modification or clearance of the compound. Amiloride hydrochloride treatment did not detectably block bacterial entry, and FK2 colocalization experiments did not reveal a clear change in bacterial ubiquitylation. These findings suggest that the amiloride-associated infection phenotype is more consistent with perturbation of the host-cell environment than with direct antibacterial activity or direct impairment of bacterial ubiquitylation resistance.

Atwal et al. reported that *O. tsutsugamushi* entry is sensitive to macropinocytosis inhibition using the amiloride derivative 5-(N-ethyl-N-isopropyl) amiloride (EIPA) (Atwal et al., 2022). Importantly, EIPA did not significantly inhibit purified OtDUB activity in our biochemical assay (Supplementary Fig. S6), indicating that the previously described effects of amiloride-related compounds on *O. tsutsugamushi* entry are mechanistically distinct from direct OtDUB inhibition. Together, these findings highlight the importance of distinguishing host-cell effects from direct inhibition of bacterial effector activity.–This distinction is particularly relevant because *O. tsutsugamushi* is an obligate intracellular bacterium and difficult to manipulate experimentally (Allen et al., 2026; Luce-Fedrow et al., 2018). Although *O. tsutsugamushi* can be propagated in mammalian cells, methods for genetically manipulating the bacterium are just beginning to be developed (Allen et al., 2026). In this setting, chemical approaches can provide useful tools to examine host-cell conditions and bacterial effector-associated phenotypes during infection, provided that direct bacterial effects and host-dependent effects are carefully separated.

Overall, this work demonstrates that OtDUB, despite its complex mode of ubiquitin recognition, can be chemically inhibited under defined experimental conditions. Identification of gentisic acid and amiloride hydrochloride shows that pathogen-encoded deubiquitylating enzymes are accessible to small-molecule modulation and provides an initial framework for developing chemical probes. Although these compounds are not optimized for OtDUB inhibition, their reproducible activity establishes the feasibility of chemical inhibition for basic investigations of *Orientia* infection mechanisms and potentially as drugs for the treatment of scrub typhus. This work also builds on reports that *O. tsutsugamushi* evades antibacterial autophagy (Choi et al., 2013; Ko et al., 2013) by confirming that it escapes ubiquitylation even in the presence of OtDUB inhibition. Hence, selective pressure of the bacterium’s intracytosolic lifestyle has likely necessitated that it evolve redundant mechanisms to counter ubiquitylation and downstream cell-autonomous immune responses.

Importantly, this study highlights the value of chemical approaches for functional characterization of bacterial effectors in systems where genetic manipulation remains limited. *O. tsutsugamushi* is an obligate intracellular pathogen for which genetic tools are just beginning to become available (Allen et al., 2026), restricting direct assessment of bacterial protein function during infection. In this setting, chemical perturbation offers one of the few practical strategies to probe effector activity in host cells. The observation that partial inhibition of OtDUB activity is sufficient to impact infection-associated phenotypes supports use of small molecules as tools to study such host–pathogen interactions. Notably, the OtDUB polypeptide includes downstream domains that catalyze guanine nucleotide exchange on the Rac1 and Cdc42 Rho-family GTPases, bind specific clathrin adaptor-protein complexes, or bind the lipid phosphatidylserine (Adcox et al., 2022; Berk et al., 2022; Lim et al., 2020). Chemical-biology approaches to interfere with these activities could also be used to dissect the functions of these OtDUB protein domains in bacterial infections of mites and mammals.

## Materials and methods

### Plasmids

All plasmids used in this study are listed in Supplementary Table 2. Plasmids were confirmed by sequencing (Berk et al., 2020; Lim et al., 2020).

### Protein purifications

DNA encoding OtDUB_1–259_ was cloned into a modified pET-28a vector encoding an N-terminal hexahistidine tag and TEV protease cleavage site, as described previously (Berk et al., 2020). Protein expression was carried out in Rosetta (DE3) *E. coli* cells. Overnight cultures grown in LB medium were back-diluted into Terrific Broth supplemented with 50 μg/ml kanamycin and grown at 37 °C to an OD_600_ of 0.6–0.8. Expression was induced with 300 μM IPTG, and cultures were incubated for 16 h at 18 °C. Cells were harvested by centrifugation and resuspended in lysis buffer containing 50 mM Tris-HCl pH 8.0, 500 mM NaCl, and 0.1 mM TCEP, supplemented with 1 mM PMSF and cOmplete protease inhibitor cocktail (Roche). Cells were lysed, and insoluble material was removed by centrifugation.

Clarified lysates were incubated with Ni-NTA agarose resin (Qiagen) for 60 min at 4 °C, washed extensively with lysis buffer, and eluted with lysis buffer supplemented with 250 mM imidazole. Eluted fractions containing OtDUB_1–259_ were pooled and concentrated, then incubated with His-tagged TEV protease at a 1:100 (w/w) ratio to remove the N-terminal His tag. TEV cleavage was performed during dialysis into ion exchange buffer (50 mM Tris-HCl pH 8.0, 0.1 mM TCEP) for 72 h. Following TEV-mediated tag cleavage, OtDUB_1–259_ was further purified by immobilized metal affinity chromatography using Ni-NTA agarose resin to remove uncleaved protein, free His tag, and His-tagged TEV protease. Final purification was performed by size exclusion chromatography using a HiLoad Superdex 75 column (Cytiva) equilibrated in 25 mM Tris-HCl pH 7.5, 100 mM NaCl, and 0.1 mM TCEP on an ÄKTA chromatography system. Fractions corresponding to monomeric OtDUB_1–259_ were pooled, concentrated, flash frozen in liquid nitrogen, and stored at −80 °C until use.

The DUB domain of CidB (Beckmann et al., 2017) and the C-terminal catalytic domain of yeast Ulp1 (Ulp1-C275) (S.-J. Li & Hochstrasser, 2003) were expressed as N-terminally GST-tagged fusion proteins using the pGEX6P-1 vector. Rosetta (DE3) *E. coli* cells transformed with the corresponding expression constructs were grown in Luria–Bertani (LB) medium supplemented with ampicillin and induced with IPTG at an OD600 of 0.5–0.6. Cultures were incubated at 18 °C overnight to allow protein expression. Cells were harvested and resuspended in ice-cold PBS containing 400 mM KCl, 1 mM DTT, and protease inhibitors. Cell lysis was performed by sonication, and clarified lysates were loaded onto a glutathione-agarose resin (Thermo) equilibrated in wash buffer (PBS with 400 mM KCl). After washing with 25 column volumes of wash buffer, GST-tagged proteins were eluted using 250 mM Tris-HCl pH 8.0, 400 mM KCl, and 10 mM reduced glutathione. Where indicated, the GST tag was removed by incubation with HRV 3C protease during dialysis. Cleaved proteins were separated from GST and protease by re-binding to glutathione resin, and the unbound fractions were collected. Final purification was performed by size exclusion chromatography using a Superdex 75 column equilibrated in 50 mM HEPES pH 7.5 and 100 mM NaCl. Peak fractions were analyzed by SDS–PAGE, pooled, concentrated, flash frozen in liquid nitrogen, and stored at −80 °C until use in enzymatic assays.

### High-throughput screening

Inhibitor screening was carried out using the Gen-Plus library (MicroSource), which consists of 960 structurally diverse, biologically active compounds, including FDA-approved drugs and compounds with known bioactivity. The library was provided by the Yale Center for Molecular Discovery and arrayed in three black 384-well plates. Compounds were dispensed into assay plates using an automated liquid-handling system at a volume of 40 nL per well. Assays were performed in AMC-cleavage buffer containing 50 mM Tris-HCl pH 7.5, 500 µM EDTA, 5 mM DTT, and 0.1% BSA, resulting in a final compound concentration of 20 µM. Vehicle control wells contained the corresponding concentration of DMSO. Purified OtDUB_1–259_ was diluted in the same AMC-cleavage buffer and added to each well at a final concentration of 125 pM. Enzyme and compounds were incubated for 60 min at room temperature before reactions were initiated by addition of ubiquitin–AMC substrate. Fluorescence was monitored over time using excitation at 345 nm and emission at 445 nm. Z′ factor values were calculated for each plate to assess assay performance. Across three screening plates, assay performance was consistent, with Z′ factor values ranging from 0.73 to 0.91.

### Compounds

Gentisic acid (Sigma-Aldrich, 149357), amiloride hydrochloride (Sigma-Aldrich, A7410), scopoletin (Sigma-Aldrich, S2500), harmol hydrochloride (Sigma-Aldrich, 116556), 5-(N-ethyl-N-isopropyl)-amiloride (EIPA; Sigma-Aldrich, A3085), and zoniporide (Sigma-Aldrich, SML0076) were dissolved in DMSO to prepare stock solutions and stored according to the manufacturer’s recommendations.

### Ubiquitin-AMC hydrolysis assays

AMC-cleavage experiments were performed as previously described (Berk et al., 2020; Shrestha et al., 2014). Ubiquitin-AMC, and SUMO1-AMC (Boston Biochem) were diluted in AMC-cleavage buffer (50 mM Tris-HCl, pH 7.5, 500 µM EDTA, 5 mM DTT, 0.1 % (w/v) bovine serum albumin), added (60 µl of 666 nM substrate) to a 96-well black polystyrene plate (Costar), and equilibrated by shaking at 30 °C for 5 min in a fluorescence plate reader (Synergy MX, BioTek). OtDUB_1–259_ diluted to the appropriate concentration was quickly added (40 µl of 8.75 nM or 875 pM enzyme) to each well, mixed for 15 s by shaking, and then datapoints were taken at 30 °C every 40 s for 60 min by 345/445 nm excitation/emission.

### Kinetic analysis of OtDUB inhibitors

Enzyme kinetic assays were performed to characterize the mode of OtDUB inhibition by gentisic acid and amiloride hydrochloride. Initial reaction velocities were measured using the fluorogenic ubiquitin–AMC substrate across a range of substrate concentrations. Reactions were carried out under the same assay conditions used above for activity and inhibition assays. Reaction measurements were obtained at increasing substrate concentrations in the absence of inhibitor and in the presence of gentisic acid or amiloride hydrochloride at concentrations of 1, 3, and 5 µM. Fluorescence was monitored continuously using excitation at 345 nm and emission at 445 nm, and initial velocities were calculated from the linear portion of the fluorescence increase. All kinetic measurements were performed using three technical replicates per condition. Lineweaver–Burk plots were generated by plotting reciprocal velocity (1/V) against reciprocal substrate concentration (1/[S]) to visualize changes in apparent kinetic parameters.

### Cell lines and pathogen cultivation

HeLa cells (CCL-2: ATCC) were cultured at 37 °C, 5% CO_2_, in Dulbecco’s Modified Eagle’s Medium (DMEM: [Gibco]) supplemented with 10% (v/v) fetal bovine serum (FBS: [Gemini]) and 1% (v/v) penicillin/streptomycin (Gibco). *Macaca mulatta* RF/6A choroidal endothelial cells (CRL-1780; ATCC) were cultured 37 °C, 5% CO_2_, in DMEM with 10% (v/v) FBS, 20 mM HEPES, and 1X non-essential amino acids (Gibco). *O. tsutsugamushi* strain Ikeda was maintained in HeLa cells as previously described (Adcox et al., 2024). *A. phagocytophilum* was cultivated in RF/6A cells and host cell-free bacteria recovered for infection experiments as previously described (Chiarelli et al., 2025).

### Transfection

For transient transfections, cells were plated in 12-well plates containing coverslips (16004-300, VWR) (1 x 10^5^ cells in 1 ml media) or in 10-cm plates (2.5 x10^6^ cells in 10 ml media). Twenty-four h later, cells were transfected using XtremeGENE9 (Roche) with pcDNA3.1(+)-based plasmids encoding full-length WT OtDUB-Flag, full-length catalytically inactive OtDUB C135A with an N-terminal 3xFLAG-6xHis tag, or the corresponding 3xFLAG-6xHis empty vector control. For this, 75 µl Opti-MEM, 3 µL XtremeGENE9, and 1 µg DNA for each milliliter of media were mixed and incubated at room temperature for 15 min before being adding dropwise to cells in antibiotic-free media.

### HeLa cell lysate preparation

HeLa cells were washed once with cold PBS and lysed directly in wells using Passive Lysis Buffer (Promega, WI) supplemented with complete protease inhibitor cocktail (Roche). Plates were incubated on ice for 20 minutes with gentle rocking. Lysates were cleared by centrifugation at 14000 × g for 20 minutes at 4 °C, and supernatants were collected. Protein concentration was determined with a Pierce BCA protein assay kit (Thermo Scientific) according to the manufacturer’s protocol, and lysate amounts were normalized prior to activity measurements.

### Cell viability assay

Cell viability was determined by CellTiter 96 AQueous One Solution Cell Proliferation assay (Promega) according to the manufacturer’s protocol. Briefly, cells were seeded at a density of 10,000 per well in 96-well plates and allowed 24 hours to attach. After cells were treated with the indicated concentrations of compounds for 72 hours, cell viability was measured using the reagent provided in the assay kit. Absorbance at 490 nm was measured using a microplate reader (Synergy MX, BioTek). Results were analyzed using GraphPad Prism.

### Infection assays

*O.* t*sutsugamushi* str. Ikeda infection assays were performed using HeLa cells as a mammalian host cell model. All experiments involving live *O. tsutsugamushi* were conducted under biosafety level 3 (BSL-3) conditions in accordance with institutional guidelines. For bacterial pre-treatment experiments, *O. tsutsugamushi* was incubated with gentisic acid or amiloride hydrochloride at final concentrations of 100 µM for 1 h at 37 °C. Following compound treatment, bacteria were washed a minimum of three times with phosphate-buffered saline (PBS) to remove residual compounds and used immediately for infection. HeLa cells were infected with treated bacteria at a multiplicity of infection of 10, confirmed as previously described (Allen et al., 2025), for 2 h at 37 °C. After infection, extracellular bacteria were removed by washing with PBS, and fresh culture medium without compounds was added. Infected cells were harvested at 2, 24, 48, and 72 h post-infection for downstream analysis.

For the host cell pre-treatment dose-response experiment, HeLa cells were treated with gentisic acid or amiloride hydrochloride at concentrations of 25, 50, or 100 µM for 48 h at 37 °C prior to infection. For other *O. tsutsugamushi* infection experiments, HeLa cells were treated with gentisic acid or amiloride hydrochloride at 100 µM for 2 or 48 h at 37 °C prior to infection. For *O. tsutsugamushi* entry and ubiquitination experiments, HeLa cells were treated with gentisic acid or amiloride hydrochloride at 100 µM for 2 h at 37 °C prior to infection. Cells were then infected with *O. tsutsugamushi* at an MOI of 10 for 2 h in the continued presence of drug. After infection, bacteria were removed by washing, and fresh medium containing the corresponding compounds was added back to the cells. Samples were collected at 2 h post-infection (hpi) for assessing *Orientia* entry and at 30 m and 2 hpi assessing for ubiquitylation of *Orientia*. For assessing ubiquitylation of *A. phagocytophilum* vacuoles, RF/6A cells were synchronously infected at an MOI of 5, determined as described previously (Chiarelli et al., 2025). At 24 hpi, the cells were fixed for downstream processing.

### Quantification of *O. tsutsugamushi* infection levels

Genomic DNA was isolated from infected cells at the indicated time points using the DNEasy Blood and Tissue Kit (Qiagen). *O. tsutsugamushi* genetic equivalents (GE) was quantified by qPCR analysis using primers specific to *O. tsutsugamushi* strain Ikeda *tsa56* gene (OTT_RS04590) (Sanchez et al., 2024) with PerfeCTa SYBR Green Fastmix (Quantbio, Beverly, MA) and a CFX384 Real Time PCR Detection System (Bio-Rad Laboratories, Hercules, CA) (Sanchez et al., 2024). Thermal cycling conditions were 95 °C for 30 s followed by 40 cycles of 95 °C for 10 s to 54 °C for 10 s, and a 65 °C-95 °C melt curve. *O. tsutsugamushi* GE ml^-1^ was calculated as previously described (Sanchez et al., 2024).

### Immunofluorescence microscopy

HeLa or RF/6A cells were grown to confluency on 12-mm glass coverslips (Bellco Glass) in polystyrene 24-well plates, followed by synchronous infection with *O. tsutsugamushi* or *A. phagocytophilum*, respectively. Samples were fixed in 4% paraformaldehyde (PFA) (32%; Electron Microscopy Sciences) diluted in 1X phosphate-buffered saline (PBS) at room temperature for 20 min, followed by permeabilization in 100% methanol for 1 min. Samples were rinsed thrice in 1X PBS, blocked in 1X PBS with 5% bovine serum albumin (Fisher) and 0.05% saponin (Sigma), followed by overnight primary antibody/antisera incubation in blocking buffer at 4 °C. Primary antibodies and antisera used for immunofluorescence were FK2 antibody (Sigma:[04-263]), TSA56 rabbit antiserum (1:1000)(Beyer et al., 2017), and APH_0032 rabbit antiserum (1:1000) (Read et al., 2022). Samples were rinsed thrice in 1X PBS and incubated for one h with Alexa Fluor 488- conjugated chicken anti-rabbit IgG and 594- conjugated goat anti-mouse IgG (ThermoFisher Scientific) at a 1:1000 dilution in 1X PBS together with 0.1 µg mL^-1^ DAPI (ThermoFisher Scientific). Three final rinses were performed and coverslips mounted with Prolong Gold Anti-Fade reagent (ThermoFisher Scientific). Micrographs were acquired with a Leica DMi8 inverted laser-scanning confocal microscope (Leica Microsystems) equipped with a Stellaris8/Falcon system and white-light laser. A 63X/1.4 NA oil immersion objective lens was used. Ubiquitin colocalization quantification was performed on an epifluorescent Leica DMi8 equipped with a Leica EL6000 lamp and 100X/1.2NA oil immersion objective lens.

### Immunoblotting

Proteins resolved in SDS-PAGE gels were transferred to PVDF membranes (Immobilon-P; IPVH00010, Millipore), and the membranes were blocked in 5% (w/v) milk in Tris-buffered saline with 0.1% Tween-20 (TBST). After blocking, the membranes were incubated in primary antibody diluted in TBST/5% milk overnight at 4 °C. Membranes were washed and then incubated with HRP-conjugated secondary antibody (NA931V and NA934V, Cytiva) in the same blocking agent as the primary antibody for 1 h at room temperature, washed and imaged by chemiluminescence (ECL) on a ChemiDoc (Bio-Rad) imaging system. Antibodies were used as follows: rabbit anti-ubiquitin (Dako, discontinued) at 1:2000; mouse anti-Flag (F3165, Sigma) at 1:5000; and mouse anti-β-actin (643802, Biolegend) at 1:5000.

### Computational docking

AutoDock Vina (Eberhardt et al., 2021; Trott & Olson, 2010) was used to perform molecular docking of gentisic acid and amiloride hydrochloride into the crystal structure of the deubiquitylase domain of the *O. tsutsugamushi* protein OTT_1962 (PDB 6UPS). (Attempts to co-crystallize OtDUB_1-259_ with either inhibitor were unsuccessful.) Docking was carried out using a cubic search box with a side length of 30 Å centered on the catalytic pocket of the DUB domain. Ligand poses were ranked based on predicted binding affinity. For each compound, the top-ranked noncovalent binding pose located within the catalytic region was selected for further analysis. The selected docking models showed predicted binding energies within approximately 95% of the highest- scoring poses and did not involve covalent bond formation. Control docking analyses were performed using the same docking workflow against representative deubiquitylating enzymes and SUMO proteases. The structures used were M48 (PDB 2J7Q), USP2 (PDB 2HD5), USP7 (PDB 4M5X), SENP1 (PDB 2XPH), USP14 (PDB 2AYN), and RickCE (PDB 5HAM). Predicted docking energies from these control analyses are shown in Supplementary Fig. S4.

### Statistical analysis and data presentation

Statistical analyses were performed using GraphPad Prism. Data are presented as mean ± SD or mean ± SEM as indicated in the figure legends. For comparisons among multiple groups, one-way ANOVA followed by the indicated multiple-comparisons test was used. For dose-response analyses, nonlinear regression was performed in GraphPad Prism. The number of independent experiments and technical replicates is indicated in each figure legend. P values < 0.05 were considered statistically significant. Micrographs were processed and analyzed using Fiji version 2.16.0/1.54 g (Schindelin et al., 2012).

## Acknowledgements

We thank the Yale Center for Molecular Discovery for assistance with small-molecule library screening. The Center is supported in part by an NCI Cancer Center Support Grant (NIH P30 CA016359). We thank Adrian B. Mehrtash for kindly providing purified USP2cc and herpesvirus M48 deubiquitylating enzymes. This work was supported by NIH grants R35 GM136325 to M.H., R01 AI167857, R01 AI139072, and R37 AI072683 to J.A.C., and F32 AI83749 to T.J.C.

## Supplementary Figure Legends

**Supplementary Figure S1.**
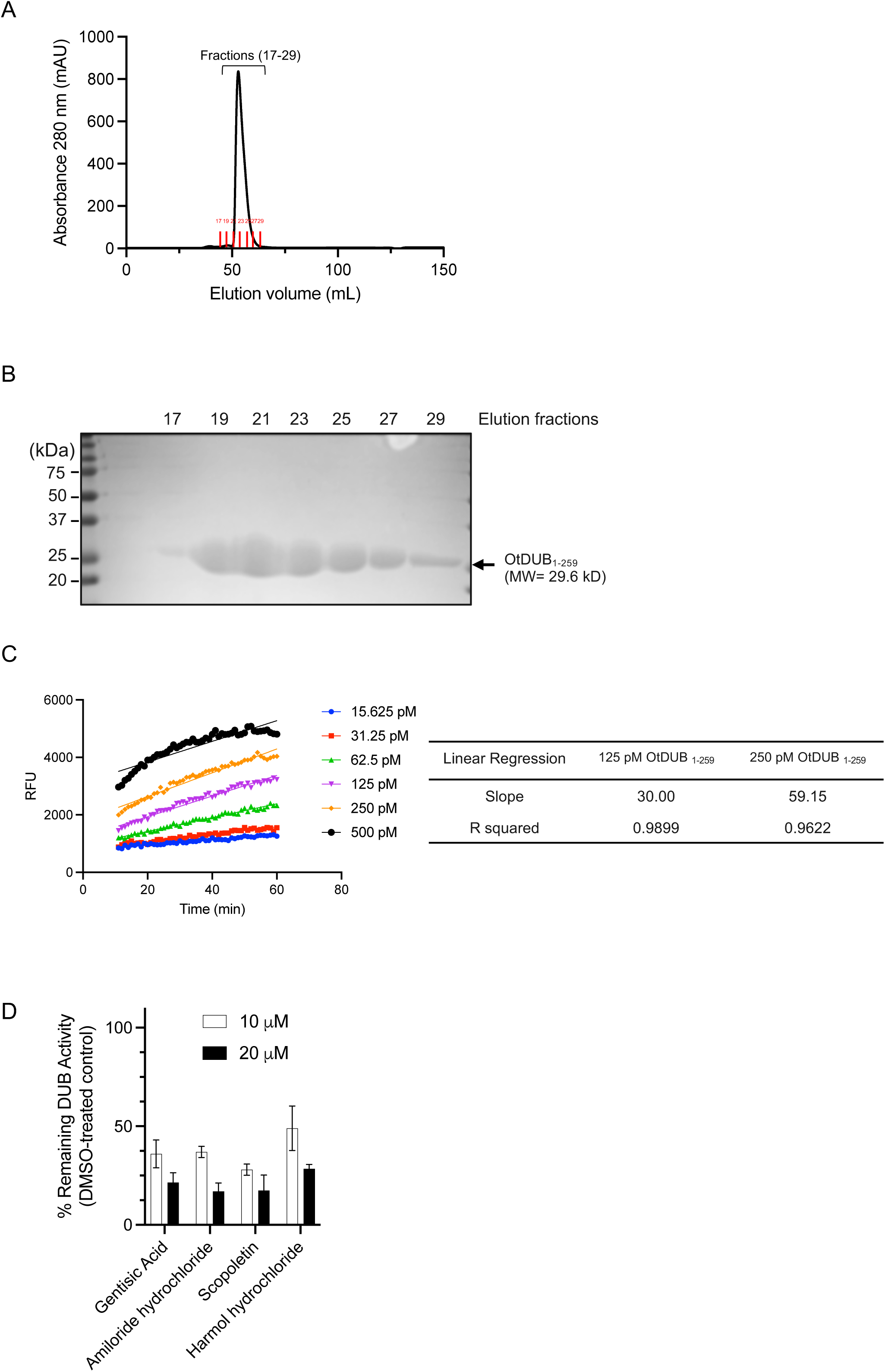
Purification and biochemical characterization of recombinant OtDUB_1–259_. (A) Recombinant OtDUB_1–259_ containing an N-terminal His tag was first purified by Ni-NTA affinity chromatography, followed by TEV cleavage to remove the His tag and final purification by Superdex 75 size-exclusion chromatography. The chromatogram shows the final size-exclusion purification step. (B) SDS–PAGE analysis of selected elution fractions from the major peak following gel filtration. OtDUB_1–259_ migrated at roughly the expected molecular weight (∼29.6 kDa). Peak fractions (19-27) were pooled for subsequent biochemical assays. (C) Time-dependent hydrolysis of the fluorogenic substrate ubiquitin–AMC by different concentrations of purified OtDUB_1–259_. Fluorescence intensity (RFU) was monitored over time to assay enzyme activity and assess linearity. Right panel summarizes representative linear regression analyses performed at the indicated enzyme concentrations. Linear increases in fluorescence signal over time confirmed that the assay conditions used for high-throughput screening were within the linear range of enzyme activity. (D) Inhibition of the truncated OtDUB_1–177_ construct by selected hit compounds. Gentisic acid, amiloride hydrochloride, scopoletin, and harmol hydrochloride were tested at 10 and 20 µM against purified OtDUB_1–177_ using the ubiquitin–AMC fluorescence assay. Residual DUB activity was normalized to DMSO-treated controls.

**Supplementary Figure S2.**
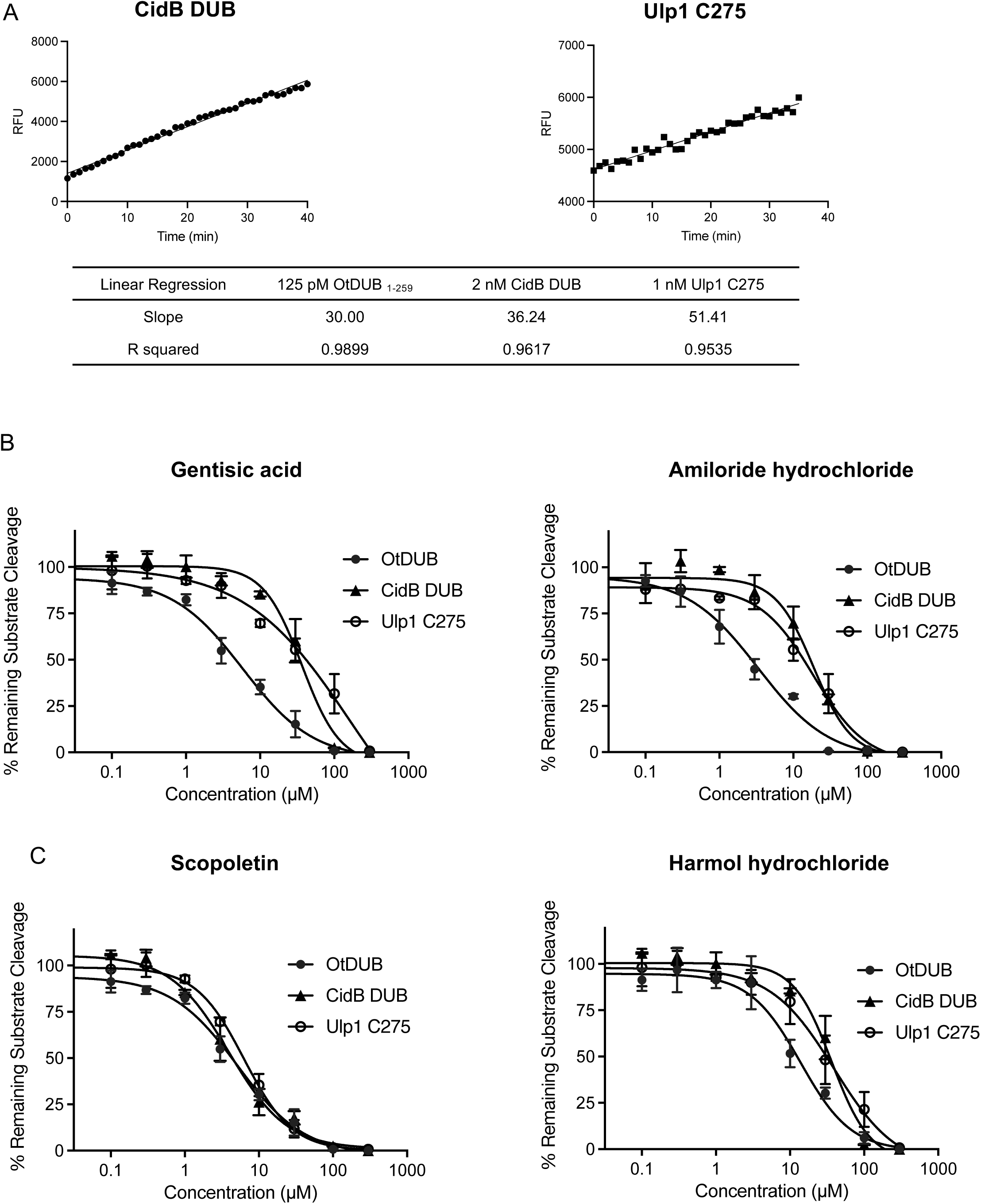
Activity characterization of the CidB*^w^*^Pip^ DUB and the Ulp1-C275 SUMO protease. (A) Time-dependent hydrolysis of fluorogenic substrates by purified CidB DUB and Ulp1-C275. Representative reaction progress curves are shown for 2 nM CidB DUB with 200 nM ubiquitin-AMC (left panel) and 1 nM Ulp1-C275 assayed with 1 µM SUMO1–AMC (right panel). Linear regression analysis was performed to compare reaction kinetics with those obtained for purified OtDUB_1–259_ (reproduced from Suppl. Fig. 1C). Linear increases in fluorescence signal over time confirmed that all enzymes were assayed within the linear range of activity under the experimental conditions used for inhibitor comparison studies. (B) Dose-dependent inhibition of purified OtDUB_1–259_, *Wolbachia* CidB DUB, and yeast Ulp1-C275 by gentisic acid and amiloride hydrochloride. Gentisic acid (left panel) and amiloride hydrochloride (right panel) were tested over the indicated concentration range with OtDUB_1–259_ and CidB DUB activity measured using ubiquitin–AMC and Ulp1-C275 activity measured using SUMO1–AMC. Residual enzymatic activity was normalized to DMSO-treated controls. (C) Dose-dependent effects of scopoletin and harmol hydrochloride on OtDUB compared with two other CE-clan cysteine proteases. Scopoletin (left) and harmol hydrochloride (right) were tested over the indicated concentration range. Reactions contained 125 pM OtDUB_1–259_ with 200 nM Ub–AMC, 2 nM CidB DUB with 200 nM Ub–AMC, or 1 nM Ulp1-C275 with 1 µM SUMO1–AMC. Residual enzymatic activity was normalized to DMSO-treated controls. DUB activity was measured relative to vehicle-treated controls under the same assay conditions.

**Supplementary Figure S3.**
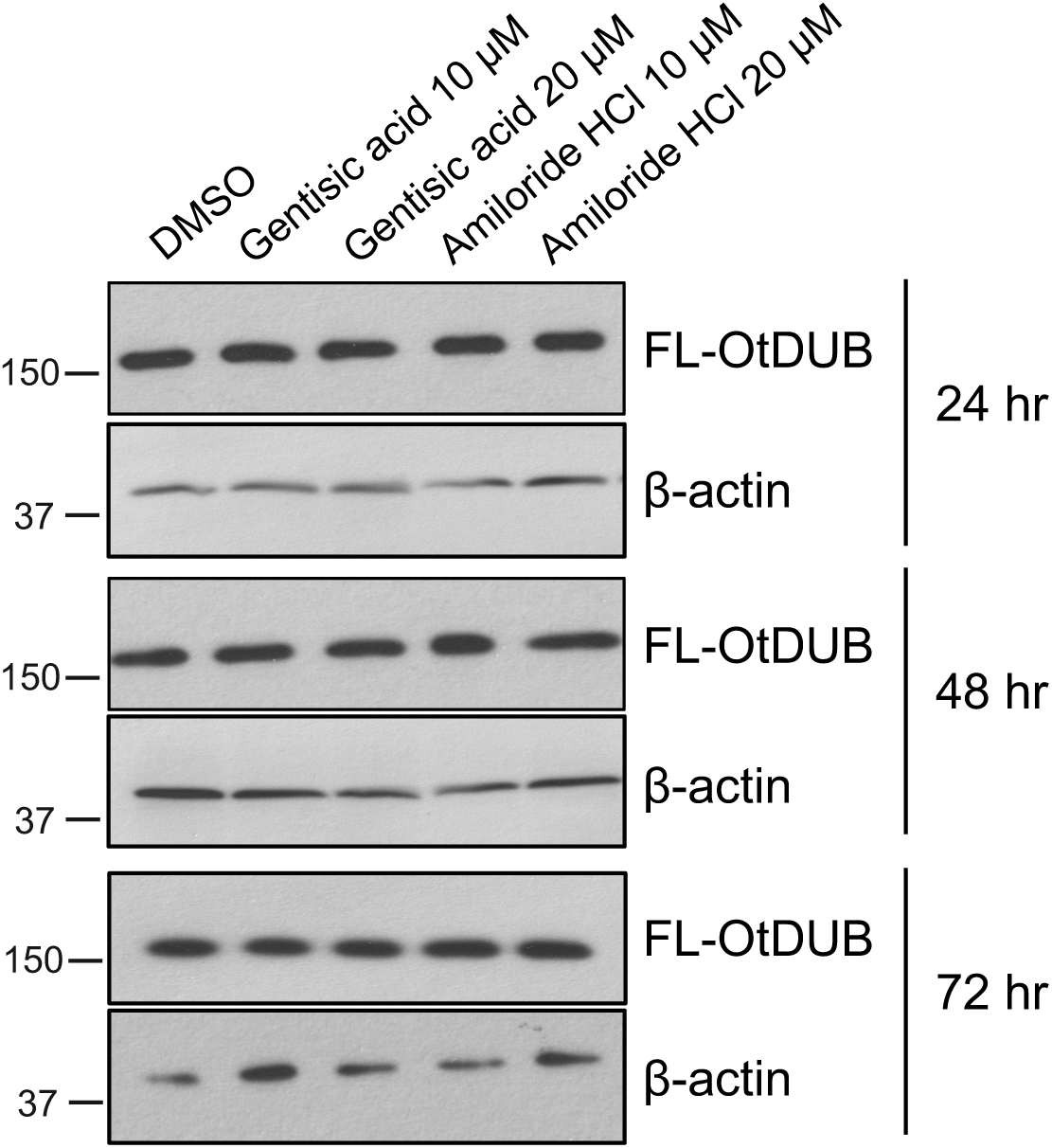
Effects of gentisic acid and amiloride hydrochloride treatment on full-length OtDUB protein levels in HeLa cells. Cells transiently expressing full-length FLAG-tagged OtDUB were treated with either compound at the indicated concentrations for 24 h, 48 h, or 72 h. Cell lysates were analyzed by anti-FLAG immunoblotting to assess OtDUB protein levels following compound treatment. β-actin was used as a loading control.

**Supplementary Figure S4.**
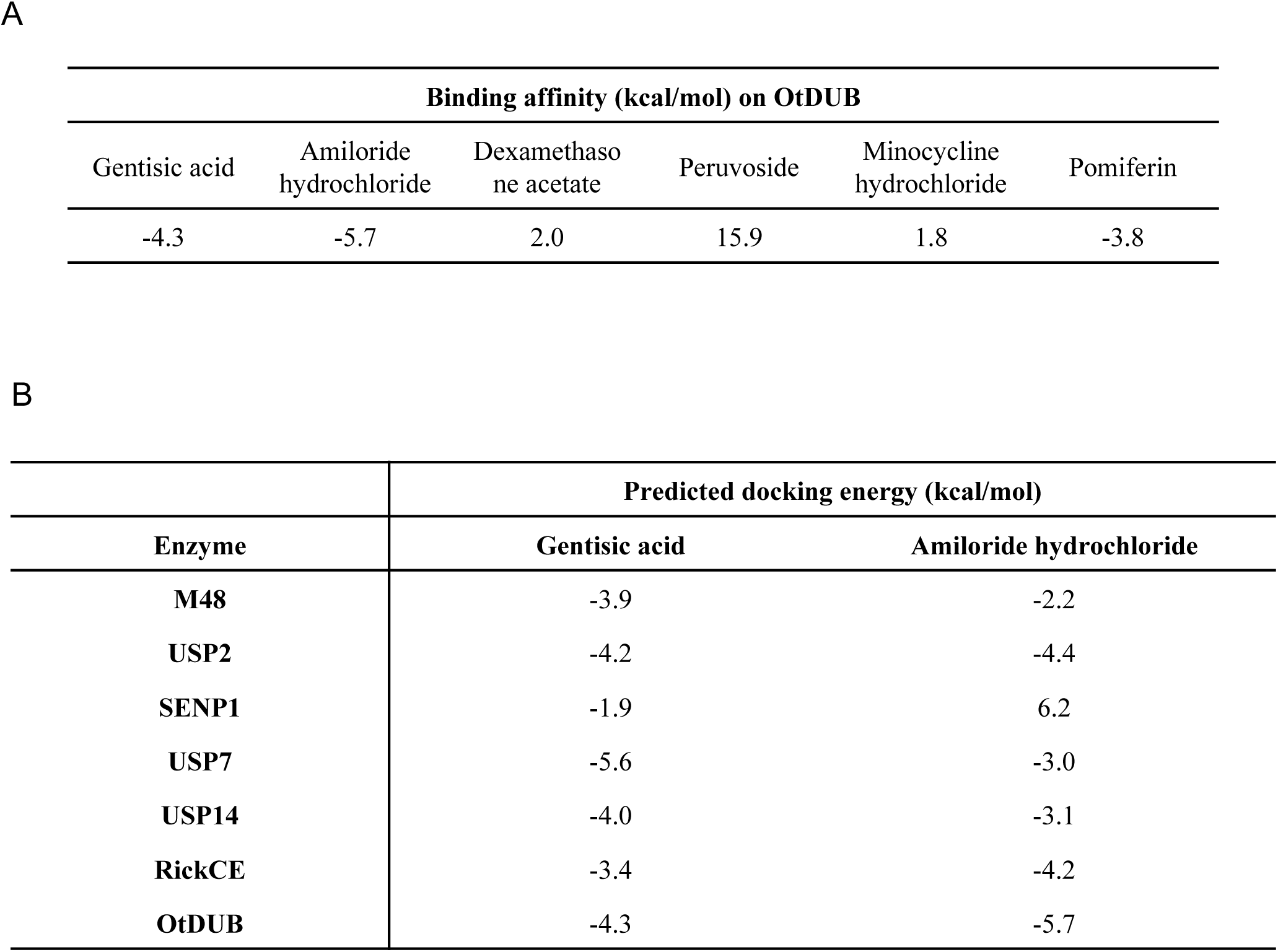
Control docking analyses using unrelated compounds and alternative deconjugating enzymes. (A) Predicted docking energies (kcal/mol) of gentisic acid, amiloride hydrochloride, and the indicated unrelated control compounds docked onto OtDUB under identical docking conditions. (B) Docking analyses of gentisic acid and amiloride hydrochloride against representative human and bacterial deubiquitylating enzymes or SUMO proteases, including M48, USP2, SENP1, USP7, USP14, and RickCE.

**Supplementary Figure S5.**
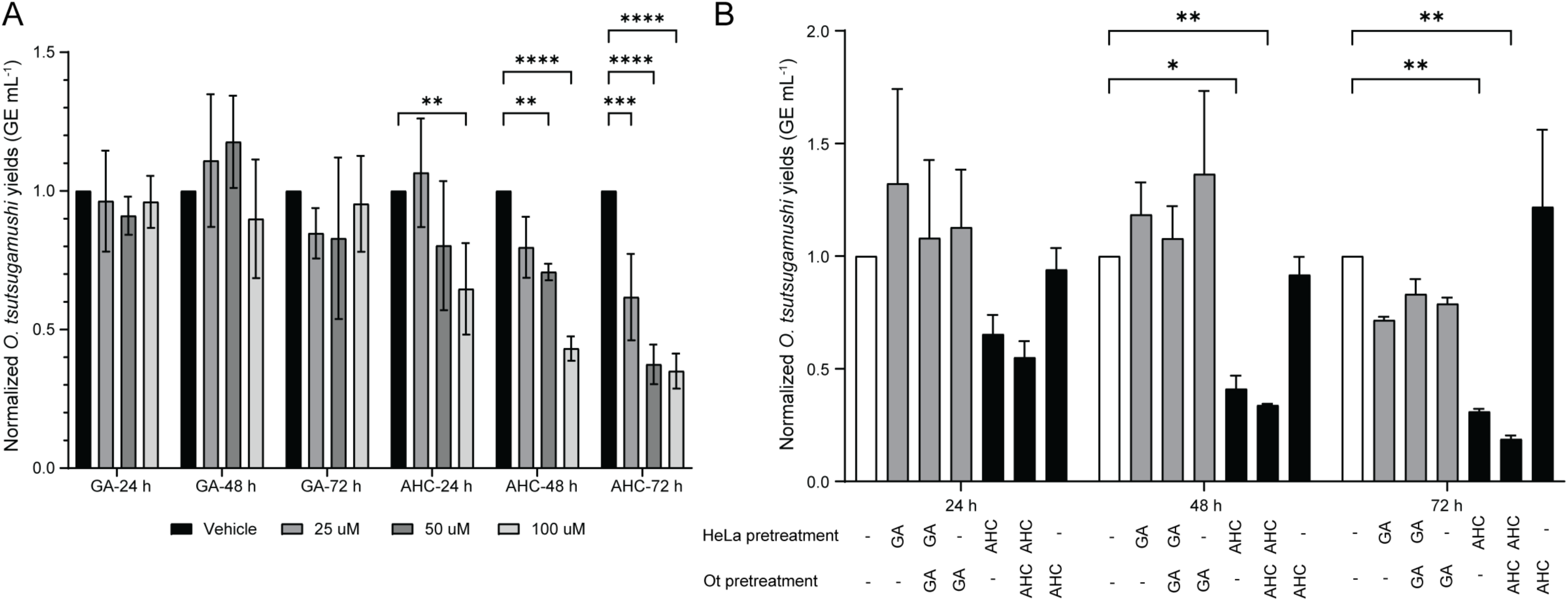
Pretreatment of host cells with amiloride hydrochloride reduces the *O. tsutsugamushi* load. (A) Effect on infection of compound pretreatment of HeLa for 48 h. HeLa cells were treated with gentisic acid (GA) or amiloride hydrochloride (AHC) at 25, 50, or 100 µM for 48 h at 37 °C followed by incubation with *O. tsutsugamushi* bacteria for 2 h. After infection, the cells were washed, fresh medium containing compound was added back to the cells, and incubation at 37 °C continued for 24, 48, or 72 h. The *O. tsutsugamushi* load in GE ml^-1^ was quantified by qPCR. (B) Comparison of the effect of compound pretreatment of host versus *O. tsutsugamushi* on infection. Host cell-free *O. tsutsugamushi* bacteria were treated with GA or AHC at 100 µM for 1 h, washed with PBS, and incubated with HeLa cells for 2 h. Alternatively, HeLa cells were treated with either compound for 2 h followed by incubation with treated or untreated *O. tsutsugamushi* organisms for 2 h. Following infection, extracellular bacteria were removed by washing and the cells were maintained in compound-free medium. Bacterial burden was quantified by qPCR. Data represent mean ± SD from three biological replicates, each measured in triplicate. Statistical comparisons were performed by two-way ANOVA followed by Dunnett’s multiple-comparisons test using infected vehicle-treated cells as the reference control. **, *P* < 0.01; ***, *P* = 0.0005; ****, *P* < 0.0001.

**Supplementary Figure S6.**
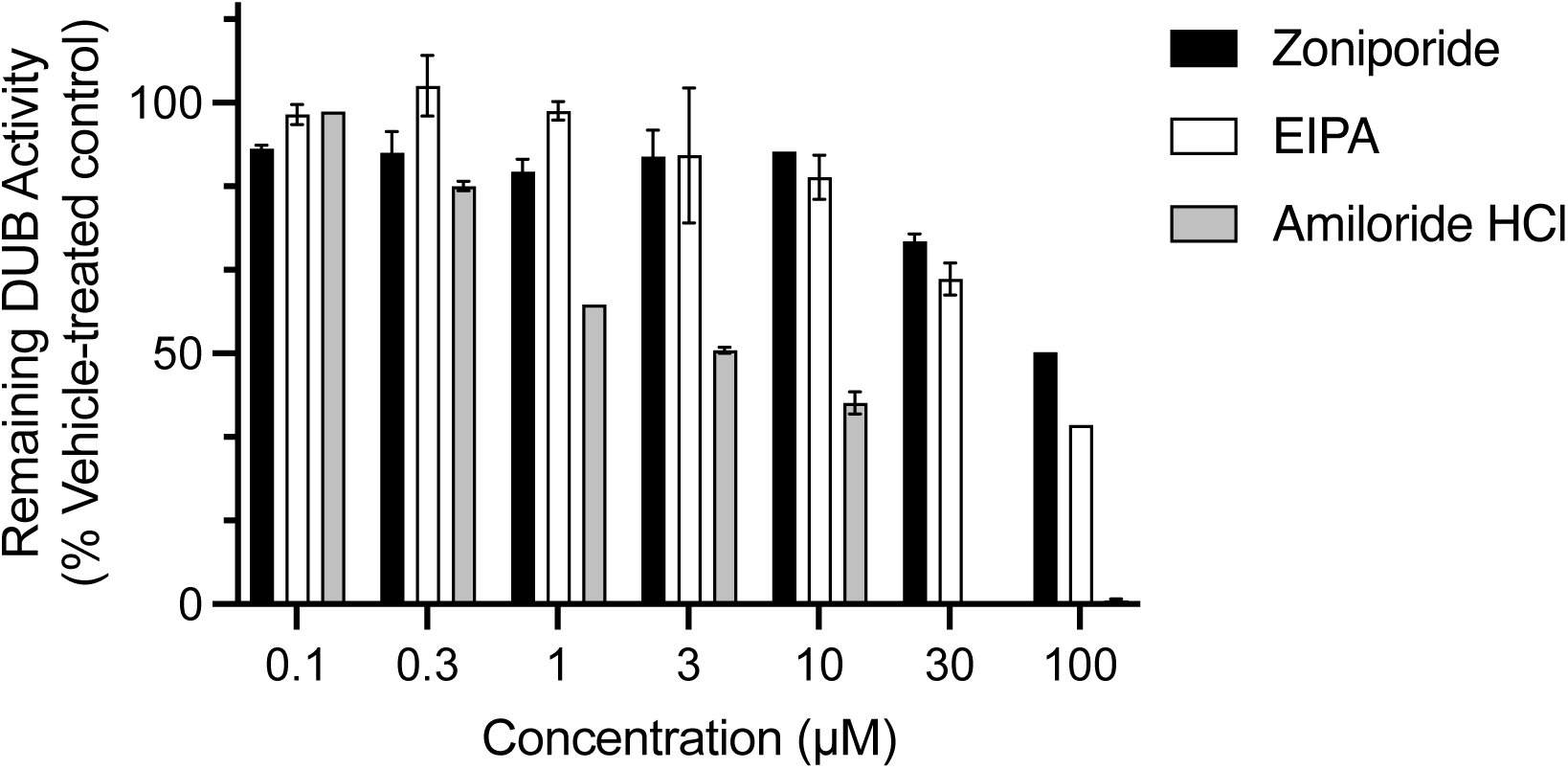
Effects of amiloride hydrochloride, EIPA, and zoniporide on purified OtDUB activity. Purified OtDUB_1–259_ was incubated with the indicated concentrations of amiloride hydrochloride, the amiloride derivative 5-(N-ethyl-N-isopropyl)-amiloride (EIPA), or zoniporide. Reactions contained 125 pM OtDUB_1–259_ and 200 nM ubiquitin–AMC, and DUB activity was measured by the AMC fluorescence. EIPA and zoniporide were included because these compounds were previously used as macropinocytosis inhibitors in studies of *O. tsutsugamushi* entry (Atwal et al., 2022). Residual DUB activity was normalized to DMSO-treated controls.

---

**Supplementary Table S1.**
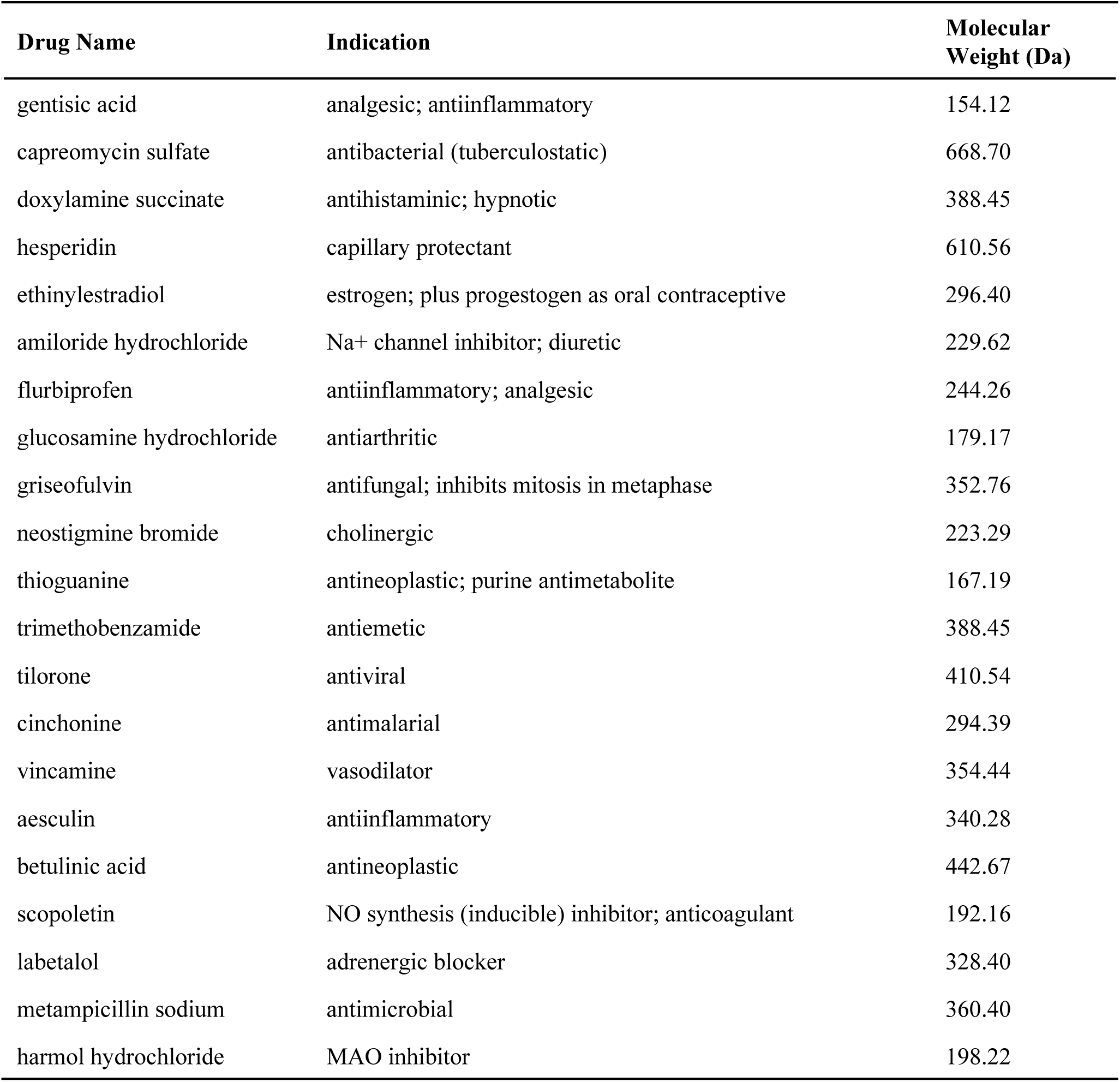

**Supplementary Table S2.**

| Plasmid | Note |
| --- | --- |
| pGEX6P1 GST-OtDUB 1-177 | (Berk et al., 2020) |
| pET28a 6His-OtDUB 1-259 | (Berk et al. 2020) |
| pCDNA3.1(+) 3xFlag-6His-empty | previously generated in MH laboratory by J. Berk |
| pCDNA3.1(+) 3xFlag-6His-Scrub DUB FL | previously generated in MH laboratory by J. Berk |
| pCDNA3.1(+) 3xFlag-6His-Scrub DUB C135A FL | previously generated in MH laboratory by J. Berk |

